# Reduced light access promotes hypocotyl growth via autophagy-mediated recycling

**DOI:** 10.1101/2021.06.16.448628

**Authors:** Yetkin Çaka Ince, Anne Sophie Fiorucci, Martine Trevisan, Vinicius Costa Galvão, Johanna Krahmer, Leonore Wigger, Sylvain Pradervand, Laeticia Fouillen, Pierre Van Delft, Sebastien Mongrand, Hector Gallart-Ayala, Julijana Ivanisevic, Christian Fankhauser

## Abstract

Plant growth ultimately depends on fixed carbon, thus the available light for photosynthesis. Due to canopy light absorption properties, vegetative shade combines reduced light and a low red to far-red ratio (LRFR). In shade-avoiding plants, these two conditions independently promote growth adaptations to enhance light access. However, how these conditions, differing in photosynthetically-available light, similarly promote growth remains unknown. Here, we show that Arabidopsis seedlings adjust metabolism according to light conditions to supply resources for hypocotyl growth enhancement. Transcriptome analyses indicate that reduced light induces starvation responses, suggesting a switch to a catabolic state to promote growth. Accordingly, reduced light promotes autophagy. In contrast, LRFR promotes anabolism including biosynthesis of plasma-membrane sterols downstream of PHYTOCHROME-INTERACTING FACTORs (PIFs) acting in hypocotyls. Furthermore, sterol biosynthesis and autophagy are indispensable for shade-induced hypocotyl growth. We conclude that vegetative shade enhances hypocotyl growth by combining autophagy-mediated recycling and promotion of specific anabolic processes.

**HIGHLIGHTS:**

- Reduced light and LRFR induce catabolism and anabolism, respectively
- Reduced light promotes autophagy to enhance hypocotyl growth in vegetative shade
- LRFR enhances hypocotyl growth by promoting plasma membrane lipid biosynthesis
- In LRFR, PIFs promote sterol biosynthesis specifically in the hypocotyl

## INTRODUCTION

Plants use a portion of electromagnetic spectrum for photosynthesis that is called photosynthetically active radiation (PAR, 400-700 nm) and composed of blue (B, 400-500 nm), green (G, 500-600 nm) and red (R, 600-700 nm) light (McCree, 1971). Leaves absorb more than 90% of B and R radiation, whereas they transmit and/or reflect a most of the far-red light (FR, 700-760 nm) (Liu and van Iersel, 2021). Therefore, plants under vegetative shade receive light combining low B (LB) and FR enrichment relative to R light leading to a low R/FR ratio (LRFR) (Casal, 2013). In shade-avoiding plants, LB and LRFR independently trigger a suite of similar adaptive responses including growth of stem-like structures e.g. hypocotyls and petioles to enhance light access (Fiorucci and Fankhauser, 2017, Buti et al., 2020, Wang et al., 2020, Ballare and Pierik, 2017). Plants in dense communities also receive LRFR due to FR reflection from neighboring leaves before actual shading. This is perceived as a neighbor proximity/shade threat signal and triggers similar growth adaptation as vegetative shade prior to a decline in light resources (Casal, 2013, Fiorucci and Fankhauser, 2017). Plant growth depends on fixed carbon (Smith and Stitt, 2007), which depends on PAR including B and R light (Moraes et al., 2019, Liu and van Iersel, 2021). Therefore, how LB and LRFR with contrasting carbon resource availability promote similar growth adaptations remains unclear (Hersch et al., 2014).

While molecular mechanisms underlying hypocotyl growth promotion are relatively well understood in LRFR, they remain unclear in LB (Buti et al., 2020, Wang et al., 2020, Ballare and Pierik, 2017). LRFR inactivates phytochrome B (phyB), leading to de-repression of Phytochrome-Interacting Factors (PIFs) transcription factors (TFs) (Buti et al., 2020, Legris et al., 2019, Wang et al., 2020, Fernandez-Milmanda and Ballare, 2021). PIF7, with substantial contributions of PIF4 and PIF5, enhances auxin production in cotyledons through induction of *YUCCAs* (*YUC2*, *YUC5*, *YUC8* and *YUC9*) coding for auxin biosynthesis enzymes (Hornitschek et al., 2012, Kohnen et al., 2016, Li et al., 2012, Muller-Moule et al., 2016, Nito et al., 2015, Nozue et al., 2015). Auxin is rapidly transported to the hypocotyl where it locally induces elongation, presumably through a combined auxin and PIF transcriptional response (Keuskamp et al., 2010, Procko et al., 2014, Oh et al., 2014, Kohnen et al., 2016). Additional hormones, notably brassinosteroids (BR) are also important for LRFR-induced hypocotyl growth (Cifuentes-Esquivel et al., 2013, Kozuka et al., 2010, Kohnen et al., 2016). Indeed, the BR signaling factor BZR1, an auxin response factor ARF6 and PIF4 collectively regulate target gene expression (Oh et al., 2014). PIFs are also key TFs for LB-induced hypocotyl elongation, where PIF4 is the primary one with contributions of PIF7 and PIF5 (de Wit et al., 2016, Keller et al., 2011, Pedmale et al., 2016). The LB response is controlled by cryptochromes (cry) but how they control PIFs remains poorly understood. The cry1-PIF4 interaction represses PIF-transcriptional activity in a B light-dependent manner (Ma et al., 2016, Pedmale et al., 2016). In LB cry2 interacts with PIF4/PIF5 but the functional importance of this complex remains unclear (Ma et al., 2016, Pedmale et al., 2016). Despite auxin and BR being indispensable for hypocotyl elongation in LB, this is not apparent from the transcriptional response, which contrasts with LRFR conditions (de Wit et al., 2016, Keller et al., 2011, Keuskamp et al., 2011, Pedmale et al., 2016, Pierik et al., 2009). Despite these differences, several growth-related pathways (e.g. cell wall organization) are transcriptionally activated in LB and LRFR (Keuskamp et al., 2011, Kohnen et al., 2016, Pedmale et al., 2016). Nevertheless, the lack of spatial resolution, limits our current understanding of LB vs LRFR growth-promoting mechanisms and the role of PIFs in hypocotyls.

Hypocotyl growth occurs by cell elongation where plasma membrane extension is essential (Gendreau et al., 1997, Hepler et al., 2013, Steer and Steer, 1989). Although the PM lipid bilayer can transit between tighter or looser packing depending on several parameters, the PM is fairly rigid with a limited expansion and contraction ability (Mamode Cassim et al., 2019). Furthermore, PM curvature is low, also limiting its extension potential (Boutté and Jaillais, 2020). In rapidly elongating plant cells (e.g., pollen tubes and root hair cells), the PM grows with the deposition of new lipid material through the fusion of Golgi-derived vesicles carrying new cell wall material (Hepler et al., 2013, Steer and Steer, 1989). We previously reported that LRFR increases carbon allocation to lipids in *B. rapa* hypocotyls (de Wit et al., 2018). Furthermore, LRFR induces sterol biosynthesis gene expression in hypocotyls (Kohnen et al., 2016). Sterols compose up to 30% of PM lipids (Mamode Cassim et al., 2019). They regulate PM permeability and fluidity (Mamode Cassim et al., 2019, Valitova et al., 2016). Together with sphingolipids, sterols are enriched in PM lipid microdomains that serve as anchoring platforms for signaling and transport proteins (Yu et al., 2020, Gronnier et al., 2019). Their major structural and functional roles at the PM suggest an important function of sterols in shade-induced hypocotyl elongation.

Production of new material required for cell elongation ultimately depends on availability of carbon compounds (Verbancic et al., 2018, Smith and Stitt, 2007). LRFR does not decrease carbon fixation in *B. rapa* seedlings, as PAR remains unchanged (de Wit et al., 2018). However, reducing PAR either by decreasing B, G or R light decreases carbon fixation (Liu and van Iersel, 2021, Moraes et al., 2019). Thus, carbon fixation is expected to decrease in LB, limiting the availability of carbon compounds that are required to promote hypocotyl growth. Carbon starvation triggered by transferring plants into darkness for several days induces autophagy that recycles unessential cytoplasmic materials by vacuolar hydrolases (Chen et al., 2019, Li and Vierstra, 2012, Wang et al., 2018). The fact that LB and LRFR differ in PAR suggests that different mechanisms may be deployed to enable cell elongation in these distinct conditions.

Analyzing light-regulated gene expression in dissected hypocotyls was informative to understand hypocotyl growth regulation during de-etiolation and in LRFR (Kohnen et al., 2016, Sun et al., 2016), but we lack equivalent data for LB. Thus, we performed organ-specific gene expression to compare and contrast the mechanisms underlying hypocotyl growth promotion in LB vs LRFR. We show that in LRFR PIFs induce PM sterols biosynthesis in hypocotyls to promote growth, while in LB autophagy is important for hypocotyl growth enhancement. Finally, we provide evidence for the need of combined LRFR-induced anabolic activities and LB-induced autophagy to enable hypocotyl elongation in vegetative shade.

## RESULTS

### LB and LRFR induce distinct transcriptional changes in elongating hypocotyls

Consistent with previous studies (de Wit et al., 2016, Pedmale et al., 2016), LB and LRFR treatments independently induced hypocotyl elongation in a PIF and YUC dependent manner (Fig. 1A). We hypothesized a convergence of LB and LRFR transcriptome in hypocotyls where both light conditions trigger cell elongation. Thus, we performed transcriptome analyses in dissected cotyledons and hypocotyls of Col-0 (wild type –WT), *pif457*, and *yuc2589* seedlings in white light (WL), LB, and LRFR to characterize the organ-specific LB and LRFR responses and their dependency on PIFs and YUC-mediated auxin biosynthesis (Fig. 1B).

**Figure 1.**
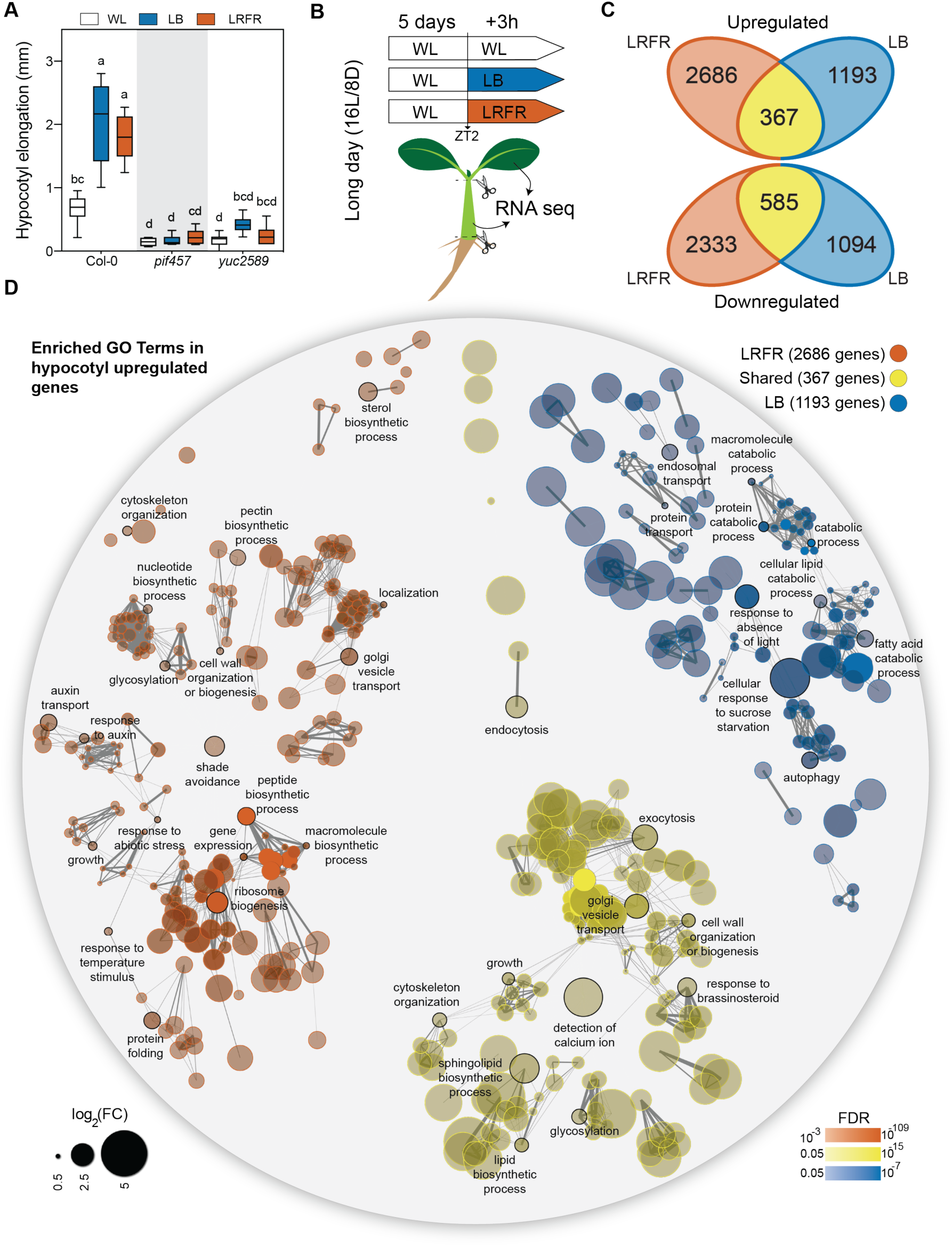
LB and LRFR induce distinct transcriptional changes in elongating hypocotyls. (A) Hypocotyl elongation of the indicated genotypes. The horizontal bar represents the median; boxes extend from the 25th to the 75th percentile, whiskers extend to show the data range. Different letters indicate significant differences (P < 0.05, n > 12, two-way ANOVA with Tukey’s HSD test). (B) Schematic summary of experimental set-up used for transcriptome analysis. (C) Number of up- and downregulated genes in Col-0 hypocotyls in the indicated light conditions (FDR<0.05, T-test with BH correction). (D) GO term enrichment analysis in Col-0 hypocotyl upregulated gene lists. Each node indicates a significantly enriched GO term (FDR<0.05). Two terms (nodes) are connected if they share 20% or more genes. The line thickness increases with the increasing number of shared genes between two terms. Only selected GO terms (black outlines) are annotated. The full list of enriched GO terms is in Table S2, the interactive version of (D) is available here. See also Figure S1.

Principle component (PC) analysis showed that biological replicates of each genotype, organ and condition clustered closely (Fig. S1A). In WT hypocotyls, LRFR induced more transcriptome changes than LB; while in cotyledons it was the opposite (Fig. 1C, S1B, Table S1). The number of common up- and downregulated genes were higher in hypocotyls than cotyledons (Fig. 1C, S1B), suggesting a convergence of LB and LRFR transcriptome in elongating hypocotyls. This was confirmed by GO term enrichment analyses for LB-specific, LRFR-specific and LB and LRFR shared upregulated genes (Fig 1D, S1C, Table S2). We highlighted selected terms for each organs and light conditions that we could easily relate to growth regulation (Fig 1D, S1C) (full lists are available in Table S2). LB and LRFR shared upregulated genes in hypocotyls were enriched in terms related to cellular elongation such as “growth”, “cell wall organization or biogenesis”, “exocytosis”, “endocytosis”, “cytoskeleton organization”, “lipid biosynthetic process” and “response to brassinosteroid” (Fig. 1D). Interestingly, genes specifically upregulated by LB in both organs were enriched in GO terms related to starvation (e.g. “cellular response to sucrose starvation”) and catabolic events (e.g. “protein catabolic process”, “cellular lipid catabolic process” and “autophagy”) (Fig. 1D, S1C). In contrast, LRFR specific genes in hypocotyls were enriched in biosynthetic processes including “ribosome biogenesis”, “peptide biosynthetic process”, “sterol biosynthetic process”, and “cell wall organization and biogenesis” (Fig. 1D). In line with previous reports (Kohnen et al., 2016), LRFR induced many hormone related responses in cotyledons and “response to auxin” in hypocotyls (Fig. 1D, S1C). Altogether, GO term enrichment analyses from light-specific and shared gene sets indicate that LB and LRFR transcriptome responses converge in elongating hypocotyls, on the induction of several growth-related mechanisms. However, we observed a striking difference between these treatments with LB upregulating numerous catabolic processes while LRFR inducing many in anabolism.

### Most LRFR-induced genes in hypocotyls require both PIFs and YUCs

In accordance with the established role of PIFs and YUCs for LRFR responses (Hornitschek et al., 2012, Kohnen et al., 2016, Li et al., 2012, Muller-Moule et al., 2016, Nito et al., 2015, Nozue et al., 2015, Lorrain et al., 2008), gene expression in *pif457* and *yuc2589* was largely unresponsive to LRFR (Fig. S1A). PCA showed that hypocotyls of LRFR-treated *pif457* and *yuc2589* grouped with WL samples but *yuc2589* cotyledons grouped closer to WT LRFR samples (Fig. S1A). We also evaluated the individual roles of PIFs and YUCCAs by analyzing the interactions between genotypes and light treatments. These comparisons revealed four groups among LRFR-upregulated genes: PIF and YUC dependent, only PIF dependent, only YUC dependent, and dependent on neither (Fig. 2A, S2A, Table S3). In hypocotyls, most LRFR-induced genes required both PIFs and YUCs, whereas the largest fraction depended on only PIFs in cotyledons (Fig. 2A, S2A). The extent of LRFR regulation was reduced in *pif457* and *yuc2589* also in PIF- and/or YUC-independent categories (Fig. 2B, S2B), indicating that our classification underestimates the importance of PIFs and YUCs. Thus, these data demonstrate that LRFR-regulated gene expression largely depends on PIF4, PIF5 and PIF7. Moreover, while in cotyledons the regulation of many genes depends on PIFs alone, in the hypocotyl their regulation depends on PIFs and YUCs.

**Figure 2.**
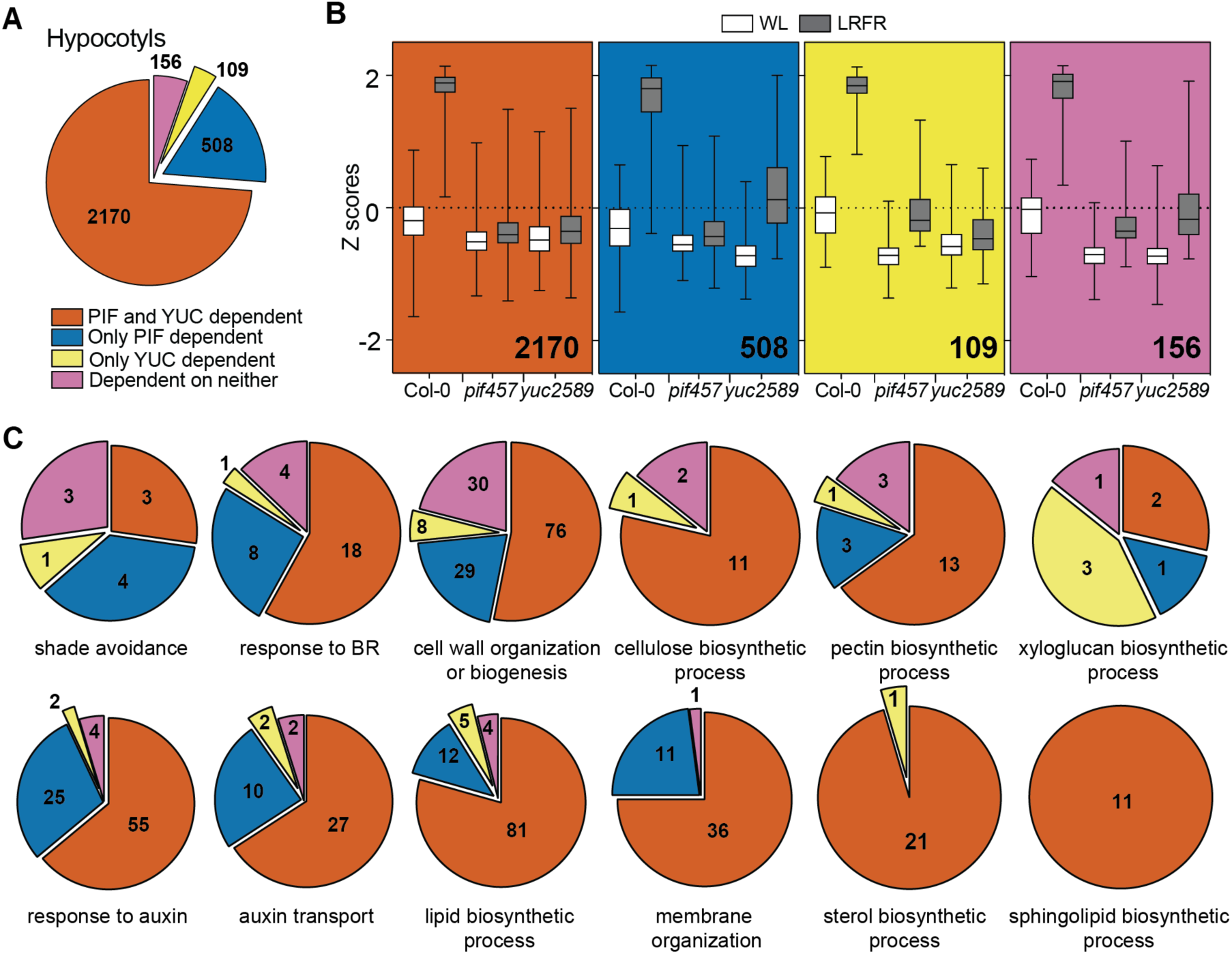
Most LRFR-induced genes in hypocotyls require both PIFs and YUCs. (A) The distribution of hypocotyl-induced genes in LRFR according to the dependence on PIFs and YUCs using the comparison of Col-0, *pif457* and *yuc2589* transcriptomes (FDR<0.05, F-test with post-hoc test). (B) Distributions of Z-scores computed from replicates averages for categories shown in (A). The horizontal bar represents the median; boxes extend from the 25th to the 75th percentile, whiskers extend to show the data range. (C) The distribution of hypocotyl-induced genes according to the dependence on PIFs and YUCs in each of the selected significantly enriched GO terms. Numbers indicate significantly regulated genes in the given categories and/or GO terms. The full list of misregulated genes and enriched GO terms are given in Table S3. See also Figure S2.

We performed GO enrichment analyses to characterize the processes depending on PIFs and YUCs (Table S3). In hypocotyls, terms such as “cell wall organization and biogenesis”, “response to brassinosteroids”, “response to auxin” and “auxin transport” heavily depended on PIFs and YUCs with a particularly strong dependency for terms related to lipid biosynthesis (Fig. 2C). In the major light-sensing organ (cotyledons), terms related to the biosynthesis of multiple hormones required PIFs but not YUCs (Fig. S2C). In contrast, genes induced in the rapidly growing hypocotyl required both PIFs and YUCs and were related to hormone responses, cell wall organization and lipid biosynthesis. Given that LRFR YUC-dependent auxin production mostly occurs in cotyledons, this indicates that in hypocotyls LRFR gene induction largely depends on auxin transported from the cotyledons with a potential local action of PIFs.

### The majority of LB-induced genes do not depend on PIFs or YUCs

Based on PCA, *pif457* and *yuc2589* displayed a robust transcriptional response to LB, contrasting with LRFR (particularly in hypocotyls) and the phenotypes of these mutants (Fig. S1A, 1A). Nevertheless, part of LB upregulated genes including some related to protein catabolism and secretion/organelle transport processes depended on PIFs and/or YUCs (Fig. 3A, 3B, S3A, list of genes and GO term analysis are given in Table S4). However, the biggest fraction of LB-induced genes did not depend on PIFs and/or YUCs (Fig. 3A, 3B, S3A). One possibility is that LB-induced hypocotyl elongation requires optimal expression of genes in WL (baseline conditions), where we found hundreds with lower expression in *pif457* and/or *yuc2589* compared to WT (Fig. 3C, Table S5). Similarly to LRFR, most WL-misregulated genes in hypocotyls required both PIFs and YUCs, whereas in cotyledons more required PIFs only (Fig. 3C, S3B). Although LB did not strongly induce these genes in the WT, the expression levels in the mutants were also lower in LB (Fig. 3D). Many GO terms related to growth, hormones and cell wall as well as “response to blue light” and “response to starvation” were enriched in the PIF- and YUC-dependent genes in hypocotyls (Fig. S3C, Table S5). Therefore, we conclude that in contrast to LRFR, LB-gene induction was less dependent on PIFs and YUCs.

**Figure 3.**
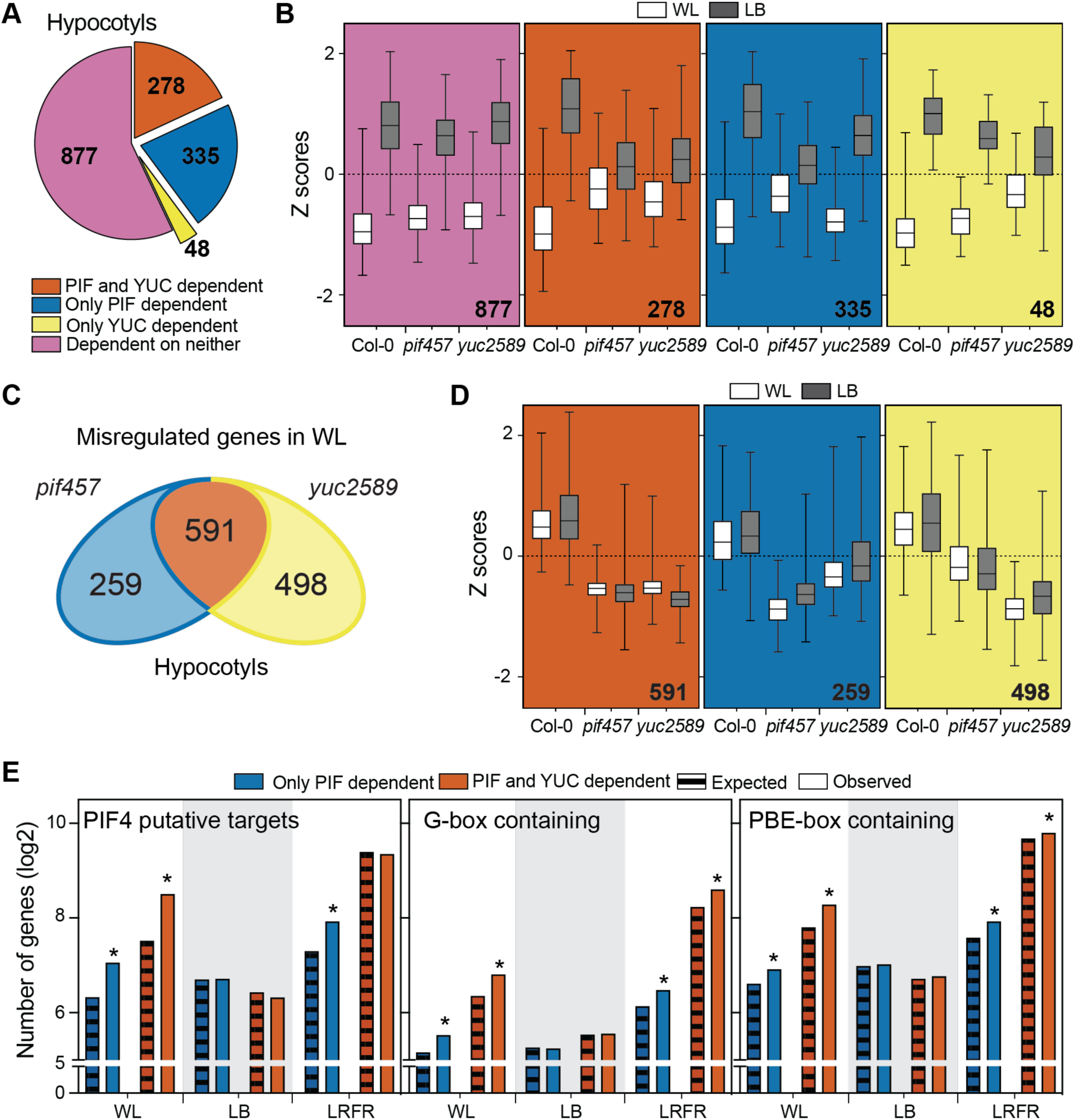
PIFs & YUCs are required for basal expression of many growth- and hormone-associated genes in hypocotyls of WL-grown seedlings. (A) The distribution of hypocotyl-induced genes in LB according to the dependence on PIFs and YUCs using the comparison of Col-0, *pif457* and *yuc2589* transcriptomes (FDR<0.05, F-test with post-hoc test). (B) Distributions of Z-scores computed from replicates averages for categories shown in (A). (C) Number of misregulated genes in *pif457* and *yuc2589* hypocotyls compared to Col-0 in WL (FDR<0.05, T-test with BH correction). (D) Distributions of Z-scores computed from replicates averages for genes that are grouped as in (C). (E) Comparison of PIF dependent genes in hypocotyls with PIF4 putative targets (as listed in (Pedmale et al., 2016)), promoters (1 kb upstream) containing G-box (CACGTG) or PBE-box (CATGTG). Asterisks (*) indicate the statistically significant overrepresentation compared to expected (P < 0.05, Binomial distribution). (B, D) The horizontal bar represents the median; boxes extend from the 25th to the 75^th^ percentile, whiskers extend to show the data range. Numbers indicate significantly regulated genes in the given categories and/or GO terms. The full list of misregulated genes, enriched GO terms, and enriched motifs are given in Table S4, S5, and S6. See also Figure S3.

Finally, we compared each set of PIF-dependent genes in WL, LB, and LRFR with PIF4 putative targets (Pedmale et al., 2016) (Table S6). PIF-dependent genes in WL and LRFR (Fig. 2, 3C, S3B) were significantly enriched in PIF4 targets for both organs (Fig. 3E, S3D). One exception was PIF and YUC dependent genes in LRFR in hypocotyls, which may indicate that these genes are induced by auxin produced downstream of PIFs (Fig. 3E). Similarly, for both organs, promoter motif enrichment analysis showed that G-box and PBE-box elements are overrepresented PIF-dependent genes in WL and LRFR but not in LB (Fig 3E, S3D, Table S6). These results suggest that PIFs directly regulate a number of genes in WL conditions, which may contribute to the impaired LB hypocotyl elongation of *pif457* mutants.

### In LRFR PIFs induce *SMT2* expression in the hypocotyl to promote growth

We confirmed that in hypocotyls LRFR induces numerous genes in diverse anabolic processes (Fig. 1D) (Kohnen et al., 2016), predominantly downstream of PIFs and YUCs (Fig. 2C, Table S3). The dependency on PIFs could be direct or indirect as they induce YUC-mediated auxin production in cotyledons (Hornitschek et al., 2012, Kohnen et al., 2016, Li et al., 2012, Muller-Moule et al., 2016, Nito et al., 2015, Nozue et al., 2015). Previously, we reported that more cotyledon-fixed carbon was allocated into the lipid fraction of elongating hypocotyl in LRFR in *B. rapa* (de Wit et al., 2018). Lipid biosynthesis, that is required for membrane expansion in rapidly elongating cells (Hepler et al., 2013, Steer and Steer, 1989), was a prominent example of anabolic processes regulated by PIFs in LRFR (Fig. 2C). We focused on sterols because of a global up-regulation of the pathway where many genes may be direct PIF targets (Fig. 1D, 2C, S4A) (Chung et al., 2020, Pedmale et al., 2016). Sterols are indispensable constituents of PM and precursors of BR growth hormones and several biosynthesis mutants are either embryo lethal or show major growth defects (Valitova et al., 2016). C-24 sterol methyltransferases (SMT) coded by two paralogs *SMT2* and *SMT3* synthesize the predominant PM sterol, sitosterol (Fig. 4A) (Carland et al., 2010, Carland, 2002, Hase et al., 2005, Mamode Cassim et al., 2019, Valitova et al., 2016). Sitosterol levels decrease dramatically in *smt2* and marginally in *smt3* mutants but these mutants still contain high levels of other sterols (e.g. campesterol) and BR and they do not have serious growth defects despite a decrease in the major PM sterol (Carland et al., 2010, Carland, 2002, Hase et al., 2005). This enabled us to conduct physiological and molecular experiments using these mutants.

**Figure 4.**
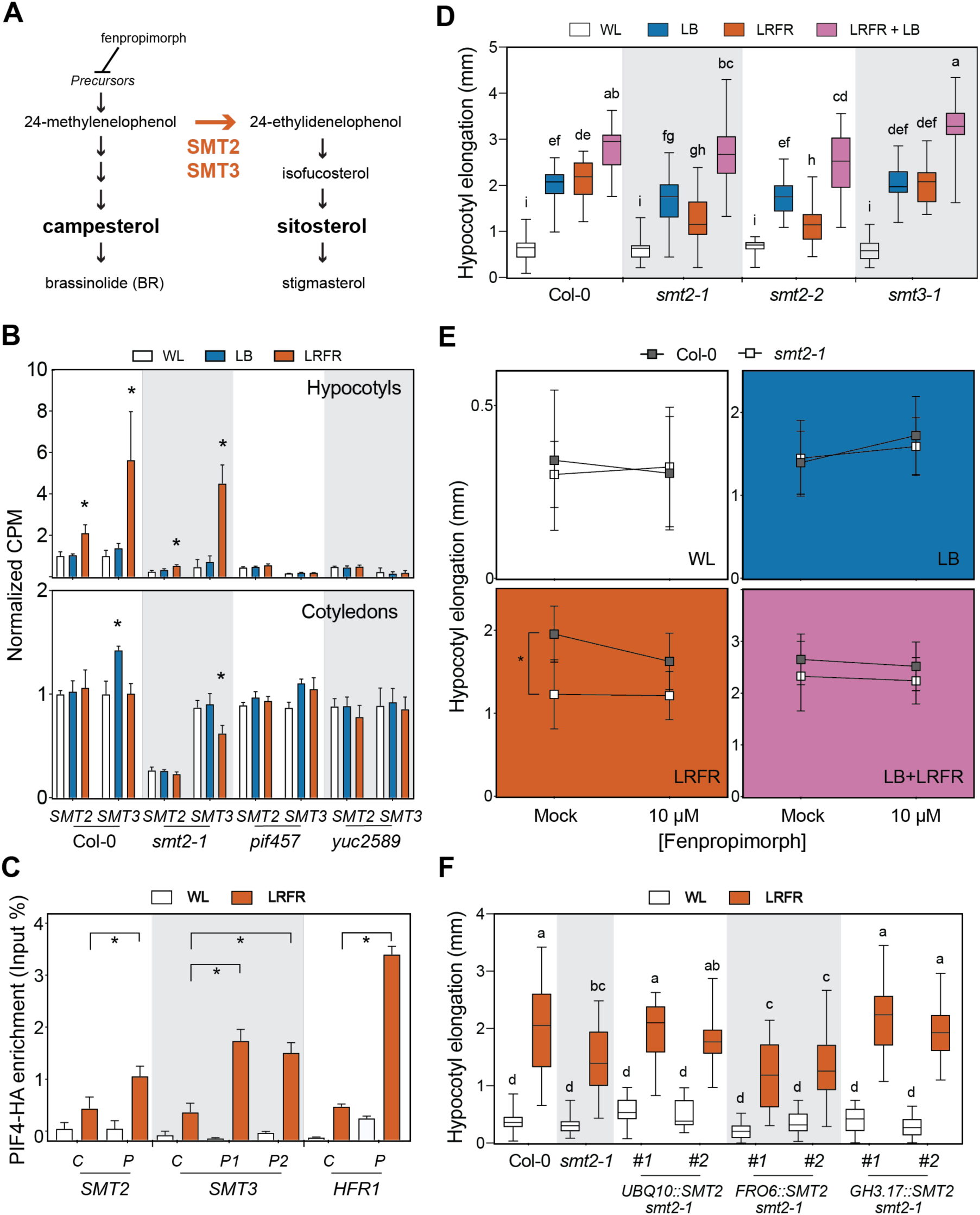
SMT2 is required locally for LRFR-induced hypocotyl elongation. (A) A simplified representation of sterol biosynthesis pathway in *Arabidopsis* (Carland et al., 2010). (B) Normalized CPM (counts per million, normalized to Col-0 average in WL) of *SMT2* and *SMT3* in the indicated genotypes and conditions (data from RNA-seq). (C) PIF4-HA binding to the promoter of the indicated genes. PIF4-HA enrichment is quantified by qPCR and presented as IP/Input (n=3, technical). Control – C, Peak – P. (D, E, F) Hypocotyl elongation of the indicated genotypes. (D, F) The horizontal bar represents the median; boxes extend from the 25th to the 75th percentile, whiskers extend to show the data range. (B, C, E) Data are means ± SD. Different letters (D, F, two-way ANOVA with Tukey’s HSD test) and asterisks (*) (B, C, T-test) (E, two-way ANOVA) indicate significant difference (P < 0.05) compared to WL (B) or control (C), and between genotypes in given light condition (E). See also Figure S4.

*SMT2* and *SMT3* expression were induced only by LRFR selectively in the hypocotyls in a PIF and YUC dependent manner (Fig. 4B). Furthermore, LRFR led to enhanced PIF4-HA binding at the promoter regions of *SMT2* and *SMT3* (Fig. 4C) in *PIF4p::PIF4-HA* (*pif4-101)* seedlings (Zhang et al., 2017). We also detected a significant PIF7-HA enrichment on the *SMT3* but not the *SMT2* promoter (Fig. S4B) in *PIF7p::PIF7-HA* (*pif7-2)* seedlings (Galvao et al., 2019). PIF7-HA binding to *SMT3* and *HFR1*, the latter being used as a positive control, was also much lower than PIF4-HA binding (Fig. 4C, S4B). Taken together, our data demonstrate that PIFs induce *SMT2* and *SMT3* expression specifically in hypocotyls by directly binding to their promoter regions in LRFR.

Using *smt2* and *smt3* mutants (Carland et al., 2010, Carland, 2002, Hase et al., 2005), we tested the requirement of *SMT2* and *SMT3* for LRFR-induced hypocotyl elongation. Hypocotyl elongation was significantly reduced in two independent *smt2* alleles and *smt2smt3* double mutant, whereas the response to LRFR was unaffected in *smt3* (Fig. 4D, S4C). Similarly, when applied simultaneously with light treatments, fenpropimorph, that inhibit sterol biosynthesis upstream of SMTs (He et al., 2003, Mamode Cassim et al., 2019), significantly reduced the hypocotyl elongation in LRFR (Fig. 4A, 4E). Furthermore, increased drug concentration resulted in a significantly steeper reduction in hypocotyl elongation of WT compared to *smt2-1* in LRFR (Fig. S4D). Importantly, the hypocotyl elongation of *smt2* and *smt3* mutants was as in the WT in WL and LB (Fig. 4D). These phenotypes highlight the importance of sterol production during LRFR treatment, in correlation with the expression data (Fig 4B, 4D). Remarkably, LB+LRFR combination mimicking vegetative shade rescued the reduced hypocotyl elongation of *smt2-1* in LRFR (Fig. 4D). Similarly, inhibition of sterol biosynthesis simultaneously with LB and LB+LRFR did not reduce the hypocotyl elongation (Fig. 4E). This result contrasts with LB and LB+LRFR hypocotyl phenotype of *smt2smt3* that was completely impaired in sitosterol biosynthesis even before the light treatments (Fig. S4C). Collectively our results suggest that SMT2-dependent induction of sitosterol production is particularly important for LRFR-induced hypocotyl elongation.

To determine whether SMT2 is required locally for LRFR-induced hypocotyl growth, we phenotyped *smt2-1* complemented with *SMT2* coding sequence with promoters specific to cotyledon (*FRO6*) (Feng et al., 2006) or hypocotyl (*GH3.17*) (Zheng et al., 2016), in comparison to a ubiquitous (*UBQ10*) control (Fig. S4E). *UBQ10* and *GH3.17* driven *SMT2* rescued the *smt2-1* phenotype in LRFR in two independent insertion lines for each construct, whereas *FRO6* did not (Fig. 4F). Taken together our data indicate that PIF-regulated *SMT2* expression in hypocotyls is required for LRFR-induced hypocotyl elongation.

### Auxin biosynthesis, response and transport are normal in *smt2-1* in LRFR

One characteristic phenotype of the *smt2* and *smt2smt3* mutants is an impaired cotyledon vasculature pattern (cvp) (Fig. S5A) (Carland et al., 2010, Carland, 2002, Hase et al., 2005). Auxin transport from cotyledons to hypocotyls is required for LRFR-induced elongation (Keuskamp et al., 2010, Keuskamp et al., 2011, Procko et al., 2014). Thus, we determined hypocotyl growth of other severe *cvp* mutants, *cvp2* and *cvp2cvl1* (Fig. S5A) that do not interfere with sterol biosynthesis (Carland and Nelson, 2009). Interestingly, both *cvp* mutants displayed a normal hypocotyl elongation in LRFR (Fig. S5B), suggesting that the cotyledon vasculature problems of *smt2* and *smt2smt3* do not explain their hypocotyl elongation defect. Moreover, we directly analyzed auxin response in *smt2-1*. PCA and statistical analyses indicated that the transcriptome of *smt2-1* is similar to Col-0 in LB and LRFR in both organs (Fig S5C, S5D, Table S7). Transcriptional activation of the major genes coding for auxin biosynthesis in LRFR was also similar in Col-0 and *smt2-1* (Fig. 5A). Furthermore, comparison of our LRFR transcriptome and 2h picloram (synthetic auxin)-regulated genes in hypocotyls (Chapman et al., 2012) showed a high correlation between these treatments for Col-0 and *smt2-1* (Fig. 5B, Table S8), but not in *pif457* and *yuc2589* which are impaired in auxin biosynthesis in LRFR (de Wit et al., 2015, Kohnen et al., 2016, Nozue et al., 2015). The WT and *smt2-1* hypocotyls elongated similarly in LRFR with increasing doses of picloram (Fig. 5C), indicating a similar auxin response. Finally, DII-VENUS signal, a reporter for determining auxin levels (Brunoud et al., 2012), similarly decreased in hypocotyls of LRFR-treated seedlings for both genotypes, indicating increased auxin levels (Fig. 5D, 5E). All together, these results indicate that it is unlikely that the *smt2-1* hypocotyl phenotype in LRFR results from altered auxin biosynthesis, transport or response.

**Figure 5.**
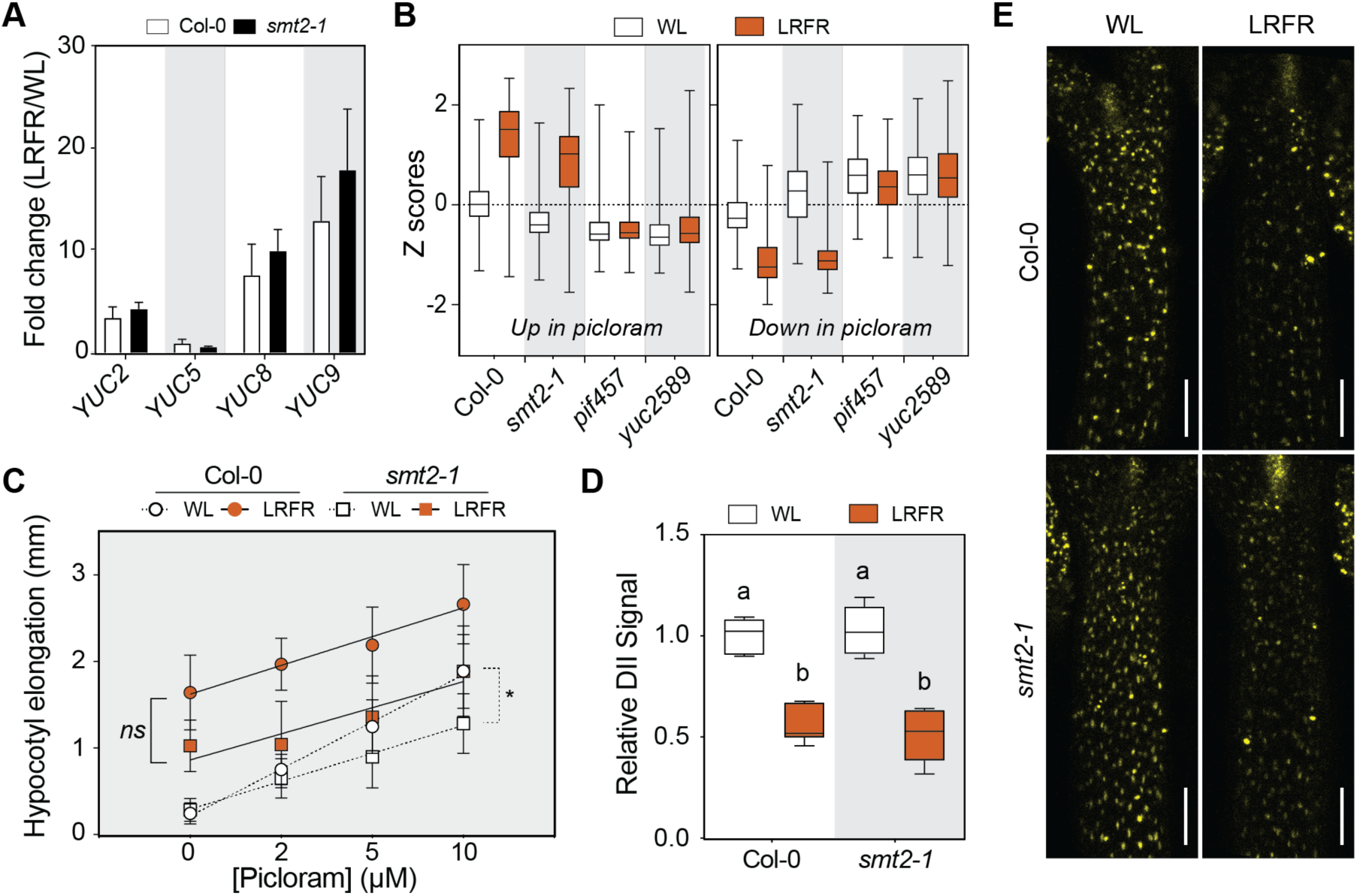
Auxin biosynthesis, response and transport are normal in *smt2-1* in LRFR. (A) LRFR-induction of auxin biosynthetic genes in the indicated genotypes (data from RNA-seq). (B) Distributions of Z-scores computed from replicates averages for synthetic auxin picloram up- and downregulated genes in hypocotyls of the indicated genotypes (as listed in (Chapman et al., 2012). (C) Hypocotyl elongation of indicated genotypes with the indicated doses of picloram (n>12). (D, E) Quantification (D) and the representative images (E) of the DII-VENUS signal intensity (normalized to mean value of Col-0 in WL) in hypocotyls of the indicated genotypes either kept at WL or transferred to LRFR for 1h (n>6). White bars equal to 100 µm. (A, C) Data are means ± SD. (B, D) The horizontal bar represents the median; boxes extend from the 25th to the 75th percentile, whiskers extend to show the data range. Different letters (D, two-way ANOVA with Tukey’s HSD test) and asterisks (*) (C, two-way ANOVA) indicate significant difference (P < 0.05) between genotypes in given light condition (C) and compared to WL (D). The full gene list of LRFR-picloram transcriptome comparison is given in Table S8. See also Figure S5.

### LRFR selectively promotes accumulation of PM lipids

We next analyzed the total lipid content globally in LRFR using untargeted lipidomics (Aldana et al., 2020) in *B. rapa* hypocotyls where LRFR enhances allocation of newly fixed carbon to the lipids and induces elongation similarly to Arabidopsis (de Wit et al., 2018). The percentage of major PM lipids (glycerolphospholipids - GPL) increased whereas the storage lipids (triacylglycerols - TAG) and the major constituents of thylakoid membranes (glycosyldiracylglycerols - GDG) (Mamode Cassim et al., 2019) decreased significantly in LRFR in the total lipid pool (Fig. 6A). We also detected a similar adjustment of lipid profile in Arabidopsis hypocotyls. We used BODIPY™ 493/503 dye that stains neutral lipids (Gocze and Freeman, 1994) to detect the level of lipid droplets (LD) that contains mainly TAG (Graham, 2008). Fluorescence intensity of LDs decreased significantly after 30h of LRFR treatment in Arabidopsis hypocotyls (Fig 6B). Furthermore, our transcriptome data showed that the “thylakoid membrane organization” GO term is enriched in the hypocotyl downregulated genes in LRFR (Table S3). In line with this data and previous reports on tomato stems (Cagnola et al., 2012), the fluorescence intensity of chloroplasts decreased in Arabidopsis hypocotyls in LRFR (Fig 6C). We conclude that LRFR promotes the accumulation of PM lipids in *B. rapa* hypocotyls whereas storage and chloroplast lipids decrease in the hypocotyls of both *B. rapa* and Arabidopsis.

**Figure 6.**
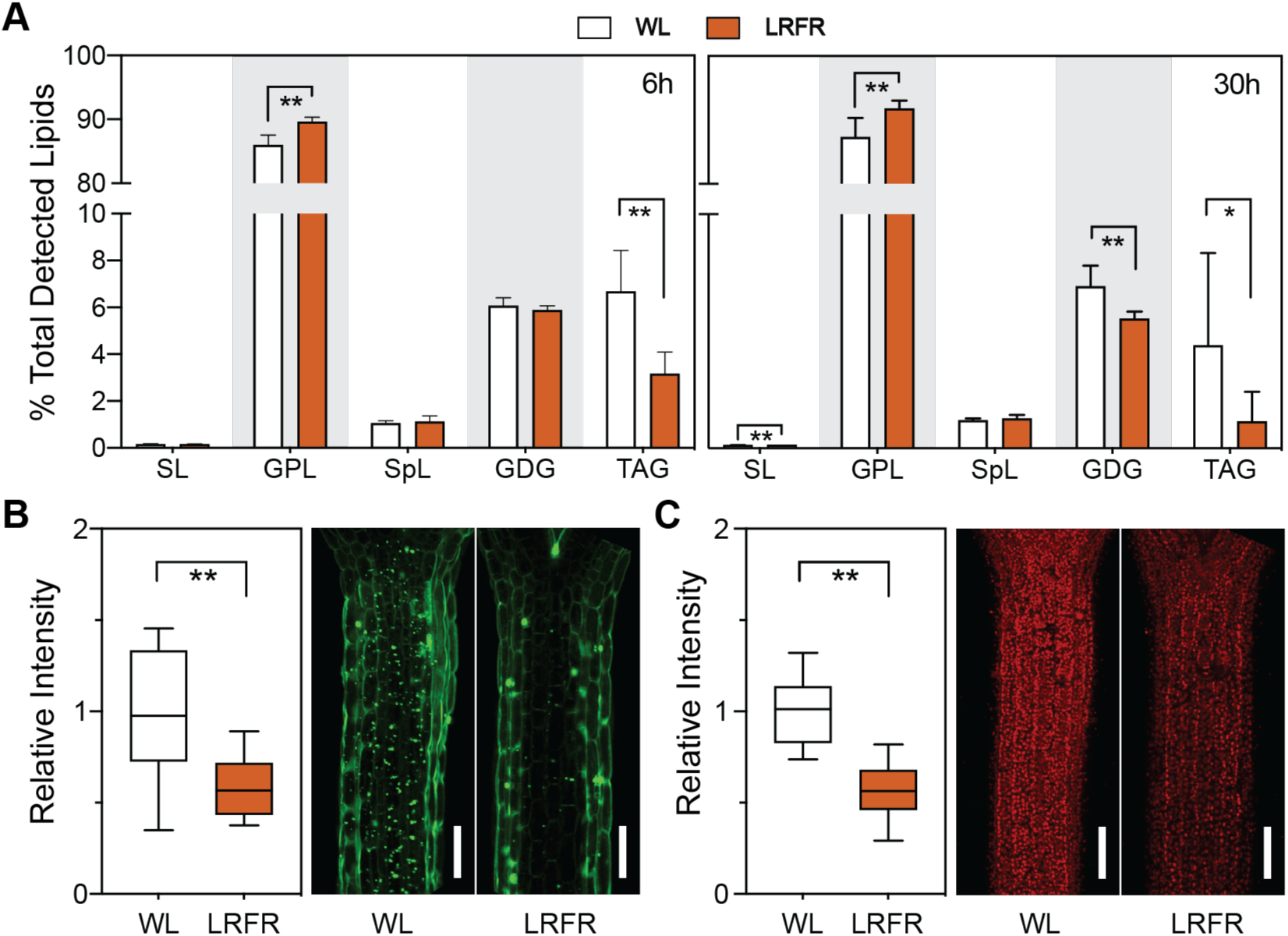
LRFR selectively promotes accumulation of PM lipids. (A) Lipid class abundance at the indicated time points is represented as percentage of total detected lipids in *B. rapa* hypocotyls (n=5, biological replicates). Sterol lipids - SL, Sphingolipids -SpL, Glycerophospholipids - GPL, Glycosyldiracylglycerols - GDG, Triacylglycerols - TAG. Data are means ± SD. (B, C) Quantification (left) and representative images (right) of (B) LDs using BODIPY™ 493/503 and (C) chloroplasts in Col-0 hypocotyls either kept at WL or transferred to LRFR for 30h (n>6). Data is normalized to WL average. White bars equals to 100 µm. (B, C) The horizontal bar represents the median; boxes extend from the 25th to the 75th percentile, whiskers extend to show the data range. (A, B, C) Asterisks indicate P values (* <0.1, **<0.05, n ≥ 5, T-test). The full list of detected lipid species is given in Table S9. See also Figure S6.

Although we focused on *SMT2* and *SMT3*, the expression of many other sterol biosynthesis genes were induced in LRFR in Arabidopsis hypocotyls (Fig. S4A) (Kohnen et al., 2016). This suggests that LRFR induces a general increase in sterols required for elongating PM. Thus, we determined the sterol composition in *B. rapa* hypocotyls where LRFR-expression profiles of *BrSMTs* were similar to their orthologs in Arabidopsis (Fig. S6A, *BrIAA29* being a control (Procko et al., 2014). Campesterol and sitosterol, the two major sterols in the PM (Valitova et al., 2016), did not change in LRFR during the timeframe of our experiments (Fig. S6B). Yet, the percentage of ergosta-5,7-dienol, a precursor for BRs downstream of campesterol, decreased after 3h of LRFR (Fig. S6B). This is in line with the report showing a decrease in another BR precursor level in LRFR (Bou-Torrent et al., 2014). These results suggest that LRFR induces a total increase in sterols rather than a major change in their composition. Taken together, our data suggest that the transcriptional upregulation of sterols and other PM lipids contribute to the buildup of PM in elongating hypocotyl cells.

### LB induces autophagy

In our LB treatment PAR was reduced to 66% of WL levels while it remained as in WL in LRFR (Fig. S7A). Such a decrease in PAR results in around 50% reduction in the net CO_2_ assimilation, regardless of the light color used for illumination (B, G, R, or their various combinations) (Liu and van Iersel, 2021). However, carbon fixation remained unchanged in *B. rapa* seedlings in LRFR (de Wit et al., 2018). In line with these observations, “carbon fixation” GO term is enriched only in LB downregulated genes, while terms related to carbon starvation responses are enriched in LB-induced genes (Fig. 1D, S7B, Table S3, S4). These data suggests that reduced light access induces a switch to a catabolic state to promote growth due to limited carbon availability. Accordingly, we observed the selective enrichment of catabolism terms and “Autophagy” in LB-induced genes (Fig. 1D, 7A). Since ATG8 is found in autophagosomal membranes (Yoshimoto et al., 2004), we used a ubiquitously expressed *mCherry*-*ATG8e* line (Hu et al., 2020) to monitor autophagic activity in our conditions. The number of autophagic bodies increased in LB-treated cotyledons when a vacuolar-type v-ATPase inhibitor, concanamycin-A (ConA) was present (Fig. 7B, S7C). We did not detect any autophagic bodies in *atg5-1,* an autophagy deficient mutant even in LB (Stephani et al., 2020, Thompson et al., 2005) (Fig. 7B). These results indicate that LB promotes autophagic flux. Further autophagic flux measurements in LB using *35S::GFP-ATG8a* (Thompson et al., 2005) demonstrated that free GFP intensity increased in LB with or without ConA, whereas the full length GFP-ATG8a decreased only in the absence of ConA (Fig. 7C, 7D). As a control, we showed that GFP band intensity remains the same in *35S::GFP* line in WL and LB (Fig. S7D). Altogether, these data indicate that LB promotes transcriptional and post-transcriptional induction of autophagy and the vacuolar degradation of autophagic bodies in Arabidopsis seedlings.

**Figure 7.**
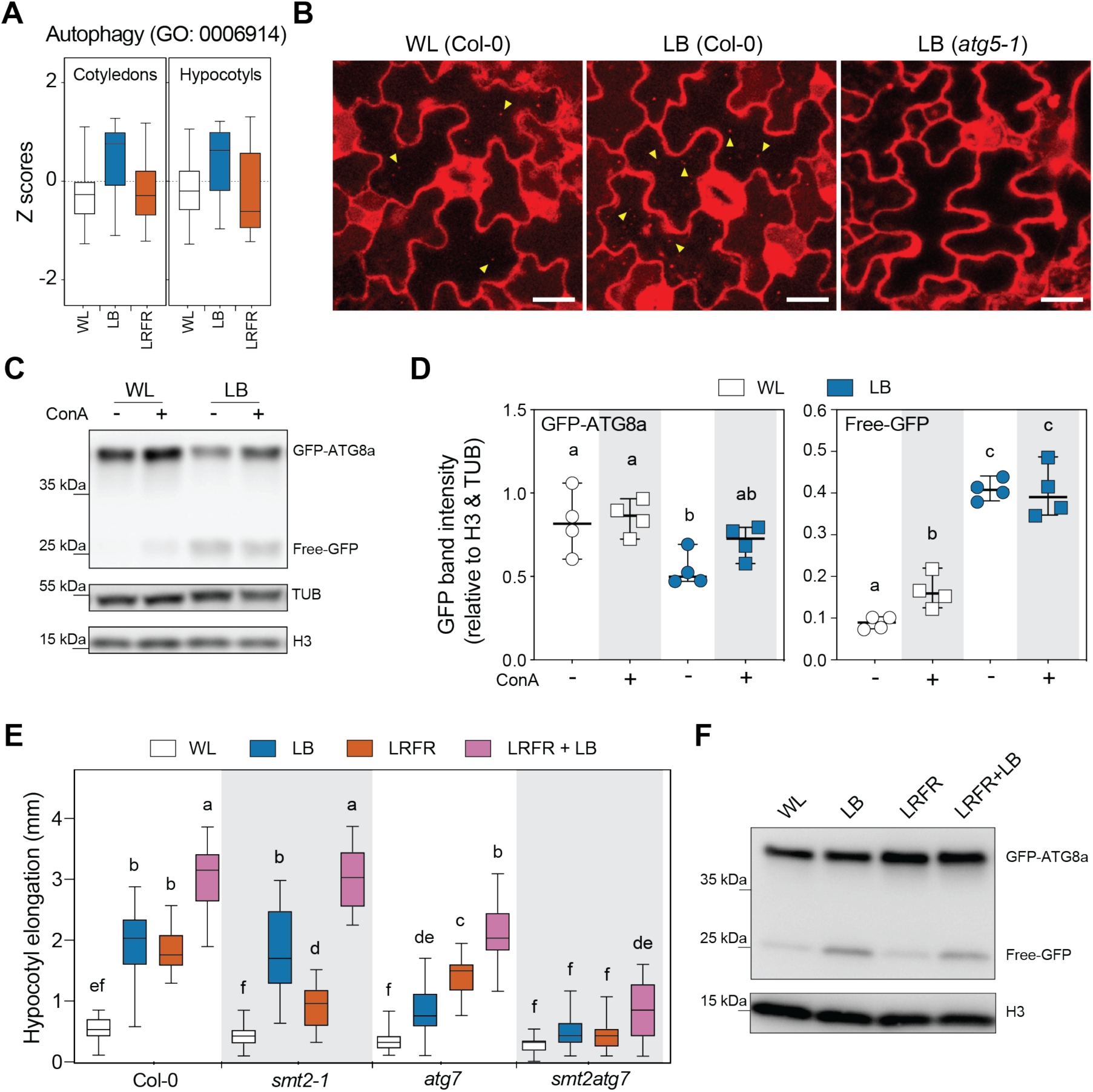
LB induces autophagy. (A) Distributions of Z-scores computed from replicates averages for genes listed in autophagy GO term in Col-0 seedlings. (B) Cotyledon pavement cells expressing *UBQ10::mCherry-ATG8e* in the indicated backgrounds either kept in WL or treated with 8h of LB in presence of concanamycin A (ConA, 5 µM). Yellow arrowheads indicate autophagic bodies. White bars equal to 20 µm. (C) Autophagic flux assay using *35S::GFP-ATG8a* (WT). GFP-ATG8a and Free-GFP levels are detected in seedlings as treated as in (B) (ConA, 0.5 µM) with an anti-GFP antibody from total protein extract. (D) Quantification of GFP-ATG8a and Free-GFP bands in autophagic flux assays (C). Each data point indicates a biological replicate (n=4 biological with average of 2 technical replicates each), horizontal bar represents the median, whiskers extend to show the data range. (E) Hypocotyl elongation of the indicated genotypes (n >12). (A, E) The horizontal bar represents the median; boxes extend from the 25th to the 75th percentile, whiskers extend to show the data range. (D, E) Different letters indicate significant difference (P < 0.05, two-way ANOVA with Tukey’s HSD test). (F) GFP-ATG8a and Free-GFP levels are detected in seedlings treated as in (C) with the indicated light conditions. H3 (C, E) and TUB (C) were used as a loading control. The full list of genes in Autophagy GO term is given in Table S10. See also Figure S7.

We next determined whether autophagy is required for LB-induced hypocotyl elongation using the autophagy mutant *atg7-2* (Hofius et al., 2009) (Fig. S7E). *atg7-2* hypocotyls elongated less in all tested light conditions except WL, with a more pronounced reduction in LB than LRFR (Fig. 7E). Remarkably, the *atg7* phenotype was complementary to *smt2-1,* which had a stronger phenotype in LRFR than LB (Fig. 7E). Moreover, LB+LRFR combination significantly enhanced hypocotyl elongation in both mutants suggesting a degree of compensation between anabolic and catabolic processes mainly promoted by LRFR and LB respectively (Fig. 7E). Supporting this idea, the *smt2atg7* hypocotyls did not elongate in either LB or LRFR (Fig. 7E). Importantly, the *smt2atg7* elongation was also very modest in LB+LRFR that induces autophagy similarly to LB (Fig. 7E, 7F, S7F). Thus, we conclude that autophagy is particularly important to promote hypocotyl elongation in LB when PAR decreases and in vegetative shade that combines LB and LRFR. Our study indicates that during the neighbor threat response LRFR-induced anabolic processes (e.g. sterol biosynthesis) fuel hypocotyl elongation, while under vegetative shade a combination of autophagy-mediated recycling and anabolic processes enable hypocotyl growth.

## DISCUSSION

In young seedlings, cotyledons are the major organs sensing LRFR while growth promotion occurs in hypocotyls (Kohnen et al., 2016, Procko et al., 2016, Procko et al., 2014). In the cotyledons, the PIF-YUC regulon controls production of auxin that promotes elongation upon transport to the hypocotyl. However, how PIFs control hypocotyl elongation locally (in the growing organ) and how PIF activity may be induced in the self-shaded hypocotyl is less clear. Modulation of auxin sensitivity in the hypocotyl appears to be one mechanism (Hersch et al., 2014, Hornitschek et al., 2012, Kohnen et al., 2016, Pucciariello et al., 2018) (Fig. 2C). Here, we show organ-specific transcriptional control of *SMT2* and *SMT3* by PIFs (Fig. 4B). ChIP data supports a direct role of PIF4 and PIF7 in the expression of those genes with enhanced binding to their promoter in LRFR (Fig. 4C, S4B). LRFR-induced *SMT2* and *SMT3* expression also depends on YUC-mediated auxin production (Fig. 4B), yet auxin alone does not induce their expression in hypocotyls (Chapman et al., 2012). This suggests a combined function of PIFs and auxin for LRFR-induced expression of *SMT2* and *SMT3*. Similarly, the vast majority of LRFR-induced genes in the hypocotyl that belong to GO categories related to growth-promoting processes depend on both PIFs and YUCs (Fig. 2, Table S3). This regulatory pattern suggests that in the self-shaded hypocotyl, an increase in auxin rather than the photoconversion of phyB may regulate PIF-mediated gene expression. This hypothesis is consistent with coordinated regulation of growth-regulatory genes by ARF6, BZR1 and PIF4 (Oh et al., 2014). Furthermore, several *PIF*s are putative targets for ARF6 and BZR1 (Oh et al., 2014). BZR1 is also known to induce *PIF4* expression during thermomorphogenesis (Ibanez et al., 2018) that induces hypocotyl elongation similarly to LRFR downstream of PIF4 and PIF7 (Chung et al., 2020, Fiorucci et al., 2019, Qiu, 2020). In contrast to hypocotyls, a substantial fraction of LRFR-induced genes in cotyledons depend on PIFs but not on YUCs (Fig. S2A). This is consistent with earlier studies identifying a number of LRFR-induced genes that do not depend on *de novo* auxin production (Tao et al., 2008). A change in light conditions in cotyledons leads to direct PIF activation potentially explaining these findings. Collectively, our data reveal organ-specific patterns of PIF-mediated gene induction in hypocotyls versus cotyledons and identify *SMT2* as an example of a gene that is selectively induced in the hypocotyl and is required for LRFR-induced elongation.

In LRFR, SMT2 promotes hypocotyl elongation locally (Fig. 4, S4). Our data suggests that this is one example illustrating the need for enhanced PM lipid production to allow hypocotyl cell elongation. LRFR leads to reallocation of newly fixed carbon to the lipid fraction of *B. rapa* hypocotyls (de Wit et al., 2018). Transcriptome analysis indicates a global upregulation of sterol and sphingolipid biosynthetic genes by LRFR (Fig. 2C). Both lipid classes are major components of the PM (Mamode Cassim et al., 2019). PM extension depends on the deposition of lipids that occurs during the delivery of membranes via exocytosis (Boutté and Jaillais, 2020, Hepler et al., 2013, Mamode Cassim et al., 2019, Steer and Steer, 1989). Of note, GO terms enriched among upregulated genes in LRFR also include endocytosis and exocytosis related terms (Fig. 1D). On the other hand, terms related to chloroplast lipids (GDG) are found among down-regulated genes in LRFR (Table S3). Consistent with the transcriptional data in Arabidopsis, PM lipids increase while chloroplast and storage lipids decrease in *B. rapa* hypocotyls (Fig. 6A). A similar decrease in chloroplasts was previously observed in tomato stems in LRFR (Cagnola et al., 2012). Moreover, we show that in Arabidopsis hypocotyls both chloroplasts and storage lipids decrease in LRFR (Fig 6B, C). Interestingly, during de-etiolation phytochromes are also known to control storage lipid utilisation (Kozuka et al., 2020). Overall, our data is consistent with LRFR leading to enhanced production of PM lipids and a reduction of other lipid classes (Fig. 6A). SMT2 and SMT3 act at a branch point of sterol biosynthesis (Fig. 4A). Their induction by LRFR may therefore also lead to changes in sterol composition which is potentially important given their role in PM fluidity and microdomain organisation (Mamode Cassim et al., 2019, Valitova et al., 2016). We currently cannot rule out this possibility, however our sterol measurements in LRFR-treated *B. rapa* do not provide strong evidence for such a change. We also note that given the global transcriptional upregulation of the sterol pathway (Fig. S4A), this is compatible with enhanced overall sterol demand but not necessarily indicative of changes in sterol composition.

PIFs and auxin production are also functionally important for LB-induced hypocotyl elongation (de Wit et al., 2016, Keller et al., 2011, Keuskamp et al., 2011, Pedmale et al., 2016) (Fig. 1A). However, the majority of LB-regulated gene expression occurred in *pif457* as in the WT (Fig. S1A). Furthermore, most of the PIF-dependent genes in LB, that represented only a small portion of LB regulated genes (Fig. 3A), are probably not direct PIF targets (Fig 3E, S3D, Table S6). Interestingly, many auxin, BR, and GA response and other growth related genes show lower basal (WL) expression in *pif457* than in the WT and this difference persists in LB (Fig. 3, Table S5). Importantly, mutant analysis and pharmacological treatments show an indispensable role for auxin, BR and GA during LB-induced hypocotyl elongation (Fig 1A) (Keller et al., 2011, Keuskamp et al., 2011, Pedmale et al., 2016, Pierik et al., 2009). We thus conclude that *pif457* gene expression in WL may contribute to the hypocotyl growth defect of the mutant in LB. This is also consistent with the report showing PIF4 and PIF5 are dose-dependent inducers of hypocotyl elongation in WL (Lorrain et al., 2008). It is however important to note that, hypocotyl elongation and PIF4 and PIF5 accumulation occurs slowly in LB compared to LRFR (Boccaccini et al., 2020, Lorrain et al., 2008, Pedmale et al., 2016, Pucciariello et al., 2018). Hence, our analysis at 3h may have missed some of the PIF-regulated transcriptional events. Therefore, the link between PIF-regulated gene expression and hypocotyl growth control in LB require further investigations.

While LRFR leads to major transcriptional changes in the hypocotyl indicative of enhanced production of many building blocks required for growth, LB leads to the induction of many catabolic processes and autophagy related genes (Fig. 1D, S1C). The LRFR gene expression pattern suggests that when light resources are fully available (Fig. S7A), the Target of Rapamycin (TOR) pathway is on and promotes e.g. the biogenesis of ribosomes and nucleotides (Brunkard, 2020). The rapid and concomitant rise in auxin and many anabolic processes in hypocotyls (Fig. 1D, S1C) (Kohnen et al., 2016) suggest a potential link between these processes. In line with this, a recent report shows that sugar-dependent TOR activity requires auxin signalling (Van Leene et al., 2019). Unlike LRFR, LB leads to PAR reduction (Fig. S7A). Reduced PAR limits carbon fixation, including in conditions where only B light is decreased (Liu and van Iersel, 2021, Moraes et al., 2019). This is in line with gene expression patterns showing LB-specific reduction in carbon fixation and induction of starvation responses (Fig. 1D, S1C, Table S3, S4). This presumably includes metabolic adjustments including alternative pathways for respiration (e.g. protein catabolism) to sustain growth in LB in line with the previous reports in other carbon limiting conditions (Araujo et al., 2010, Izumi et al., 2013, Buchanan-Wollaston et al., 2005). When energy becomes scarce, autophagy-mediated recycling is promoted (Chen et al., 2019, Goto-Yamada et al., 2019, Li and Vierstra, 2012). Consistently, LB also transcriptionally promotes autophagy and leads to enhanced production of autophagic bodies and autophagic processing of GFP-ATG8a, while LRFR does not (Fig. 7). Moreover, the *atg7* autophagy mutant had a particularly striking phenotype in LB-induced hypocotyl elongation while it was less affected in LRFR (Fig. 7E). Remarkably, this is the converse phenotype of *smt2* mutants. Interestingly, the combined light treatment, which mimics vegetative shade, largely rescues the phenotypes of both single mutants, suggesting a degree of compensation between catabolic and anabolic processes. The *smt2 atg7* double mutant phenotype with complete absence of elongation in LB and LRFR and only a modest response in combined treatments are in line with this conclusion (Fig. 7E). Importantly, *SMT2* and *SMT3* expression is induced in the hypocotyl of shade-treated seedlings (combined LB and LRFR) (Das et al., 2016). Altogether, our work indicates the requirement for enhanced *de novo* synthesis in LRFR and autophagy in LB, while the combination of both processes contributes to growth enhancement of the hypocotyl in vegetative shade.

In conclusion, our work shows that in vegetational shade that comprises a reduction in PAR and LRFR hypocotyl elongation requires autophagy and the induction of specific anabolic processes (e.g. sterol biosynthesis). In contrast, during neighbor proximity (LRFR only) growth enhancement relies on anabolic processes. We note that thermomorphogenesis and neighbor proximity both lead to similar growth adaptation using related signaling pathways (Casal and Balasubramanian, 2019, Qiu, 2020). Within a temperature range that does not significantly decrease photosynthetic efficiency, it is likely that for thermomorphogenesis as well the anabolic processes described here are relevant.

## Acknowledgements

We thank Martina Legris, Laure Allenbach Petrolati, Olivier Michaud, Mieke de Wit, Anupama Goyal, Ana Lopez Vazquez, Ganesh Mahadeo Nawkar and Maud Lagier for providing resources, technical support and/or comments on the manuscript; René Dreos for advice on statistical methods; Niko Geldner, Ersin Gül (ETH Zurich), Sinem Celebioven (University Zurich), Soner Yildiz (Icahn School of Medicine) and Yasin Dagdas (GMI, Vienna) for suggestions and comments on the manuscript; Christian Hardtke, Teva Vernoux (ENS Lyon), Richard Vierstra (Washington University, St Louis), Dany Geelen (University Gent) and Yasin Dagdas for providing resources. We are grateful to the CIF (https://cif.unil.ch/) for help with microscopy, the GTF (https://wp.unil.ch/gtf/) for RNA seq. experiments and the Metabolomics Unit (https://www.unil.ch/metabolomics/en/home.html) for the lipidomic analysis and the Bordeaux-Metabolome platform for sterol analysis (https://www.biomemb.cnrs.fr/en/lipidomic-plateform/). Work in the Fankhauser lab is supported by the University of Lausanne and the Swiss National Science Foundation (310030B_179558), the Bordeaux Metabolome Facility-MetaboHUB by a grant from ANR (no. ANR–11–INBS–0010).

## Author contributions

YCI: Conceptualization, Formal analysis, Data curation, Methodology, Investigation, Resources, Validation, Visualization, Writing – original draft. ASF, MT and VCG : Investigation, Resources, Validation. JK: Validation. SP and LW: Formal analysis, Data curation, Methodology . SM : Conceptualization, Data curation, Validation, Investigation, Methodology. LF and PVD : Investigation. JI and HGA: Investigation, Data curation, Validation. CF: Conceptualization, Data curation, Methodology, Project administration, Resources, Funding acquisition, Supervision, Validation, Writing – original draft. All authors read and approved the manuscript.

## RESOURCES AVAILABILITY

### Lead Contact

Further information and requests for resources and reagents should be directed to and will be fulfilled by the Lead Contact, Christian Fankhauser

### Materials availability

This study did not generate new unique reagents. Plasmids and transgenic plants generated in this study are available from the Lead Contact with a completed Materials Transfer Agreement.

### Data and code availability

The RNA-seq data discussed in this publication have been deposited in NCBI’s Gene Expression Omnibus (Edgar et al., 2002) and are accessible through GEO Series accession number GSE174655 (https://www.ncbi.nlm.nih.gov/geo/query/acc.cgi?acc=GSE174655).

## EXPERIMENTAL MODEL AND SUBJECT DETAILS

### Plant material and growth conditions

We used the *Arabidopsis thaliana* genotypes (cv Columbia-0) listed in key resources table. *yuc2yuc5yuc8yuc9* was recrossed using all *yuc* alleles that are described in (Nozue et al., 2015) except *yuc5-1* (SAIL_116_C0). We used the strain R-o-18 for *B. rapa* experiments. Oligonucleotides used for genotyping are listed in Table S11.

Seeds were size-selected and surface-sterilized using 70% (v/v) ethanol and 0.05% (v/v) Triton X-100 for 3 min followed by 10 min incubation in 100% (v/v) ethanol. Seeds were sowed on ½ Murashige and Skoog medium (½ MS) containing 0.8% (w/v) phytoagar (Agar-Agar, plant; Roth) and subsequently stratified at 4 °C for 3 day in darkness. For hypocotyl elongation and RNA-seq experiments where seedlings were grown on vertical plates the phytoagar concentration was raised to 1.6% (w/v) (Ince and Galvao, 2021). For all experiments, seedlings were grown in 16h/8h, light/dark photoperiod (LD) at 21 °C in a Percival Scientific Model AR-22L (Perry, IA, USA) incubator. WL was emitted from white fluorescence tubes (Lumilux cool white 18W/840) at a fluence rate of 130 μmol m^-2^s^-1^ and LRFR was achieved by supplementing WL with 45 μmol m^-2^s^-1^ FR light (LEDs) lowering the R (640–700 nm)/FR (700–760 nm) from 1.4 to 0.2, as measured by Ocean Optics USB2000+ spectrometer. A double layer of yellow filter (010 medium yellow, LEE Filters) lowering blue light from 7 μmol m^-2^s^-1^ (WL) to 0.5 μmol m^-2^s^-1^ (LB) was used to cover up the seedlings for LB treatments. The light spectra are shown in Figure S7A.

For hypocotyl elongation and microscopy experiments seedlings were grown for 4 days in WL and subsequently kept in WL or transferred to light treatment (at ZT2) for additional 3 days. Figure 1A was an exception where seedlings were grown for 5 days in WL and subsequently kept in WL or transferred to light treatment (at ZT2) for additional 3 days. Pharmacological treatments (fenpropimorph and picloram) were done on vertically grown seedlings on nylon meshes. After 4 days in WL seedlings on nylon meshes were transferred to new plates containing the drug or the corresponding solvent and put for 3 additional days into WL or light treatment (at ZT2). Fenpropimorph (Carbosynth, United Kingdom, FF23264), picloram (Sigma-Aldrich, Steinheim, Germany, P5575), and Concanamycin A (Sigma-Aldrich, Steinheim, Germany, C9705) were dissolved in DMSO (dimethylsulfoxide) applied at the indicated concentrations in Figure legends (DMSO for mock).

For RNA-seq and western blot experiments seedlings were grown for 5 days in WL and subsequently kept in WL or transferred to light treatment (at ZT2) for 3h (RNA-seq) or 8h (western blots) before harvesting. For concanamycin A treatment, seedlings were transferred to liquid ½ MS with shaking (75 rpm).

For ChIP-qPCR experiments, seedlings were grown in WL for 5 days and then either kept in WL or transferred to LRFR (at ZT2) for additional 5 days before harvesting.

For RT-qPCR, complex lipid and sterol measurement analysis, *B. rapa* seedlings were grown for 5 days in WL and subsequently kept in WL or transferred to LRFR (at ZT2) for the indicated time in legends before harvesting.

Seedlings imaging and measurements were described previously (Ince and Galvao, 2021). In short, hypocotyl length was measured from images before (day 4) and after the treatments (day 7). Hypocotyl elongation for each individual seedling is calculated by subtracting the length at 4d from the length at 7d..

## METHOD DETAILS

### Constructs cloning

PCR amplifications were performed using Phusion® High-Fidelity DNA Polymerase (New England Biolabs, Massachusetts, USA, Cat. No. M0530). All cloning was done using In Fusion® HD Cloning kit (Takara, California, USA; Cat. No. 639649). First, *GUSPlus::tOCS* was cloned in *pFP100* plasmid carrying *pAt2S3::GFP* selection marker (Bensmihen et al., 2004) and the new plasmid was named as *pYI001. pFR06* and *pGH3.17* were cloned into *pYI001* in order to obtain *pFRO6::GUSPlus::tOCS* and *pGH3.17::GUSPlus::tOCS*, respectively*. pUBQ10::SMT2-Flag::tOCS* was cloned into *pFP100,* while *pFRO6::SMT2-Flag,* and *pGH3.17::SMT2-Flag* were cloned into *pYI001.* The primers are listed in Table S11. These constructs were transformed into *smt2-1* plants using *Agrobacterium tumefaciens GV3301* strain by floral dip (Clough and Bent, 1998).

### RNA isolation, quantitative RT-PCR and RNA-sequencing

For RNA isolation, 5d-old seedlings were harvested in liquid nitrogen and kept at -70 °C for overnight. Next day, seedlings were covered with -70 °C cold RNA*later*™-ICE (Thermo Fisher Scientific, United States, AM7030) and transferred to -20 °C overnight. Cotyledons and hypocotyls were dissected using sharp needles on top of an ice block under a binocular microscope (Nikon, SMZ1500) and RNA isolation and reverse transcription quantitative polymerase chain reaction (RT-qPCR) reactions were performed as previously described (Kohnen et al., 2016). Oligonucleotides are listed in Table S11.

For RNA-sequencing, RNA quality was assessed on a Fragment Analyzer (Agilent Technologies). From 40 ng total RNA, mRNA was isolated with the NEBNext Poly(A) mRNA Magnetic Isolation Module. RNA-seq libraries were then prepared from the mRNA using the NEBNext Ultra II Directional RNA Library Prep Kit for Illumina (New England Biolabs, Massachusetts, USA). Libraries were quantified by a fluorimetric method and their quality assessed on a Fragment Analyzer (Agilent Technologies). Cluster generation was performed with the resulting libraries using Illumina HiSeq 3000/4000 SR Cluster Kit reagents. Libraries were sequenced on the Illumina HiSeq 4000 with HiSeq 3000/4000 SBS Kit reagents for 150 cycles. Sequencing data were demultiplexed with the bcl2fastq Conversion Software (v. 2.20, Illumina; San Diego, California, USA).

### ChIP-qPCR

10d-old seedlings *PIF4p::PIF4-HA* in *pif4-101* (Zhang et al., 2017) and *PIF7p::PIF7-HA* in *pif7-2* seedlings (Galvao et al., 2019) grown in WL for five days and then either kept in WL or transferred to LRFR for another five days were harvested in liquid nitrogen. Chromatin extraction was performed as described previously (Bourbousse et al., 2018) except that samples were cross-linked only with formaldehyde. Immuno-precipitation was performed as described previously (Gendrel et al., 2005) using an anti-HA antibody (Santa Cruz Biotechnology, Inc., Dallas, TX, USA; sc-7392 X). The qPCR was done in triplicates or quadruplicate on input and immunoprecipitated DNA. Peaks (P) were defined using a genome-wide ChIP study from etiolated seedlings that identified PIF4-peaks (Oh et al., 2012). Controls (C) are on coding regions of each gene. Oligonucleotides are listed in Table S11.

### Western-blot analysis

Total protein extracts from seedlings were obtained as previously described (Galvao et al., 2019). Protein samples were separated on 10% Mini-Protean TGX gels (Bio-Rad, Hercules, CA, USA) and blotted on nitrocellulose membrane (Bio-Rad) using Turbo transfer system (Bio-Rad). Membranes were blocked with 5% milk overnight at 4°C or 1h at room temperature for Anti-GFP JL-8 (1:4000; Clontech, California, USA; Cat. No. 632380/632381), polyclonal H3 (1:2000; Abcam, Cambridge, UK; Cat. No.1791), Anti-TUB (1:2000; Abicode, California, USA, Cat. No. M0267-1a) antibodies before probing with horseradish peroxidase (HRP)-conjugated anti-rabbit (for H3) or anti-mouse (for anti-GFP and anti-TUB) as the secondary antibody (1:5000; Promega, Madison, USA; Cat. No. W4011 and W4021, respectively). Chemiluminescence signal were obtained with Immobilon Western Chemiluminescent HRP Substrate (Millipore, Merck KGaA, Darmstadt, Germany) on an ImageQuant LAS 4000 mini (GE Healthcare, Buckinghamshire, UK). Images were processed with ImageJ software (http://rsb.info.nih.gov/ij).

### Microscopy and GUS Staining

DII-Venus microscopy and image quantification were performed as indicated in (Kohnen et al., 2016). In short, 4-day-old seedlings grown as described above were transferred to LRFR or kept in WL for 1h before imaging. We used an inverted Zeiss confocal microscope (LSM 710, 20X objective, 0.8 DIC). VENUS signal was detected using an Argon laser (excitation at 514 nm and band pass emission between 520 and 560 nm). Image stacks (5-6/seedling) were acquired for every hypocotyl until the VENUS signal was lost. The pinhole was opened to collect the maximal signal intensity together with the minimal stack number (5.42 airy units, 20.2 μm section, 10.08 μm interval). Images were processed with ImageJ software (http://rsb.info.nih.gov/ij). To quantify the VENUS signal, we used the SUM slices projection of 4 slices from the stack excluding the first layer with the stomata.

For the lipid droplet (LD) quantification, 4-day-old seedlings grown as described above were transferred to LRFR or kept in WL. Seedlings were treated with BODIPY ™ 493/503 (2 µg/mL in H_2_O) for 20 min at room temperature and imaged on inverted Zeiss confocal microscope (LSM 710, 20X objective, 0.8 DIC) 30h after the beginning of light treatment. BODIPY ™ 493/503 signal was detected using an Argon laser (excitation at 488 nm and band pass emission between 500 to 540 nm). Image stacks (6/seedling) were acquired for every hypocotyl. The pinhole was opened to 3.15 Airy Units (11.4 µm section, 10.00 µm interval). Images were processed with ImageJ software (http://rsb.info.nih.gov/ij). To quantify the signal, we selected a region of interest (ROI) on the upper half of the hypocotyls and analyzed the fluorescence intensity of particles (size=0-75 circularity=0.50-1.00) in each image stack with a threshold (>3000) and summed the total intensity using the “Analyze particles” tool.

For chloroplast quantification, we used the same protocol as the LD quantification with the following changes. We used a HeNe633 laser (excitation at 633 nm and band pass emission between 640 to 685 nm to have an equal contribution from ChlA and ChlB). The pinhole was opened to 3.15 Airy Units (11.4 µm section, 10.00 µm interval). Threshold was adjusted to >10000; size and circularity were kept as default (0-infinity and 0-100, respectively) in ImageJ.

For visualization of autophagic bodies in *UBQ10::mCherry-ATG8e* lines (Hu et al., 2020, Stephani et al., 2020), we used the following settings. 4-day-old seedlings grown as described above were transferred to LB or kept in WL for 8h before imaging in the presence (5 µM) or absence (DMSO) of concanamycin A (Sigma-Aldrich, Steinheim, Germany, C9705). We used DPSS 561-10 laser (excitation at 561 nm and band pass emission between 570 to 635 nm) (LSM 710, 63X objective, 1.3 oil DIC). The pinhole was opened to 5.32 Airy Units (4.3 µm section, 4.21 µm interval). 2 stacks are combined for each image.

The protocol for GUS staining reactions was described in (Galvao et al., 2019). Cotyledons were prepared for cotyledon vasculature imaging as described in (Carland, 2002). GUS staining and cotyledon vasculature were imaged using a dissecting microscope (Nikon SMZ1500).

### Sterol measurements

4 hypocotyls from 5d-old *B. rapa* seedlings per sample were pooled and frozen in liquid nitrogen immediately after fresh weights were recorded. Samples were heated for 1h in EtOH with 1% H_2_SO_4_ at 85°C. Sterols were extracted in hexane. Free hydroxyl groups were derivatized at 110°C for 30 min, surplus BSTFA-trimethylchlorosilane (Sigma-Aldrich, Steinheim, Germany, Cat. No. B-023) was evaporated and samples were dissolved in hexane for analysis using GC-MS (Agilent 7890, A coupled to a mass spectrometer, MSD 5975, Agilent EI) under the same conditions as described (Cacas et al., 2016). Quantification of sterols was based on peak areas, which were derived from total ion current and using cholestanol as internal standard. Each sterol was normalized to the total amount of detected sterols and presented as percentage of total.

### Untargeted lipidomics mass spectrometry analysis

4 hypocotyls from 5d-old *B. rapa* seedlings per sample are pooled and preheated isopropanol (at 75 °C) was added immediately after fresh weights are recorded. Each sample with isopropanol was incubated at 75 °C for 15 min to inhibit phospholipase activity and cooled down to room temperature. Samples were kept at 4 °C overnight and isopropanol was then evaporated to dryness using Nitrogen steam. Dry extracts were then reconstituted in 200 μL of IPA spiked with the internal standard mixture (SPLASH® LIPIDOMIX® Mass Spec Standard (92/8; v/v)). This solution was further homogenized in the Cryolys Precellys 24 sample Homogenizer (2 x 20 seconds at 10000 rpm, Bertin Technologies, Rockville, MD, US) with ceramic beads. The bead beater was air-cooled down at a flow rate of 110 L/min at 6 bar. Homogenized extracts were centrifuged for 15 minutes at 21000 g at 4°C (Hermle, Gosheim, Germany) and the resulting supernatant was collected and transferred to an LC-MS vial.

Extracted samples were analyzed by reversed phase liquid chromatography coupled to a high-resolution mass spectrometry (RPLC-HRMS) instrument (Agilent 6550 IonFunnel QTOF).  In both, positive and negative ionization mode, the chromatographic separation was carried out on a Zorbax Eclipse Plus C18 (1.8 μm, 100 mm × 2.1 mm I.D. column) (Agilent technologies, USA). Mobile phase was composed of A = 60:40 (v/v) Acetonitrile:water with 10 mM ammonium acetate and 0.1% acetic acid and B = 88:10:2 Isopropanol:acetonitrile:water with 10 mM ammonium acetate and 0.1% acetic acid. The linear gradient elution from 15% to 30% B was applied for 2 minutes, then from 30% to 48% B for 0.5 minutes, from 48% to 72% B and last gradient step from 72% to 99% B followed by 0.5 minutes isocratic conditions and a 3 min re-equilibration to the initial chromatographic conditions. The flow rate was 600 μL/min, column temperature 60 °C and sample injection volume 2μl.

ESI source conditions were set as follows: dry gas temperature 200 °C, nebulizer 35 psi and flow 14 L/min, sheath gas temperature 300 °C and flow 11 L/min, nozzle voltage 1000 V, and capillary voltage +/- 3500 V. Full scan acquisition mode in the mas range of 100 – 1700 *m/z* was applied for MS1 data acquisition while MS/MS data were acquired in the iterative data dependent acquisition mode to facilitate lipid identification and annotation.

Pooled QC samples (representative of the entire sample set) were analyzed periodically (every 6 samples) throughout the overall analytical run in order to assess the quality of the data, correct the signal intensity drift and remove the non-reproducible signals (CV > 25%) (Dunn et al., 2011). In addition, a series of diluted quality controls (dQC) were prepared by dilution with isopropanol: 100% QC, 50%QC, 25%QC, 12.5%QC and 6.25%QC and analyzed at the beginning and at the end of the sample batch. This QC dilution series served as a linearity filter to remove the features that do not respond linearly (correlation with dilution factor is < 0.65) (Gagnebin et al., 2017).

### Data processing

Raw LC-HRMS and HR(MS/MS) data was processed using MS-Dial software (Tsugawa et al., 2015) (http://prime.psc.riken.jp/Metabolomics_Software/MS-DIAL/). Relative quantification of lipids was based on EIC (Extracted Ion Chromatogram) areas for the monitored precursor ions at the MS1 level. Peak areas were normalized considering the sample amount (mg) (full lists of lipid species are given in Table S9). The obtained tables (containing peak areas of detected and identified lipids by MS and MS/MS, and using MS only across all samples) were exported to “R” software http://cran.r-project.org/ where the signal intensity drift correction was done within the LOWESS/Spline normalization program (Tsugawa et al., 2015) followed by noise filtering (CV (QC features) > 30%) and visual inspection of linear response.

The abundance of each MS/MS detected lipid species was normalized to the total amount of MS/MS detected lipids and presented as a percentage of total.

## QUANTIFICATION AND STATISTICAL ANALYSES

### RNA-seq initial data analysis

Purity-filtered reads were adapters and quality trimmed with Cutadapt (v. 1.8) (Martin, 2011). Reads matching to ribosomal RNA sequences were removed with fastq_screen (v. 0.11.1). Remaining reads were further filtered for low complexity with reaper (v. 15-065) (Davis et al., 2013). More than 30 million uniquely mapped reads were obtained per library and reads were aligned against the *Arabidopsis thaliana.TAIR10.39* genome using STAR (v. 2.5.3a) (Dobin et al., 2013). The number of read counts per gene locus was summarized with htseq-count (v. 0.9.1) (Anders et al., 2015) using the *Arabidopsis thaliana.TAIR10.39* gene annotation. Quality of the RNA-seq data alignment was assessed using RSeQC (v. 2.3.7) (Wang et al., 2012).

Statistical analysis was performed for genes independently in R (R version 4.0.2). All steps described here were performed separately for the samples from hypocotyls and cotyledons (except for the initial clustering of all samples together). Genes with low counts were filtered out according to the rule of 1 count(s) per million (cpm) in at least 1 sample. The number of genes retained in the analyses based on this filtering is different for hypocotyls and cotyledons. Library sizes were scaled using TMM normalization. Subsequently, the normalized counts were transformed to cpm values and a log2 transformation was applied by means of the function cpm with the parameter setting prior.counts = 1 (edgeR) (Robinson et al., 2010).

Differential expression was computed with the R Bioconductor package “limma” (Ritchie et al., 2015) by fitting data to a linear model. The approach limma-trend was used. Fold changes were computed and a moderated t-test was applied. P-values were adjusted using the Benjamini-Hochberg (BH) method, which controls for the false discovery rate (FDR), globally across several comparisons of experimental conditions. The adjustment was performed in a few different ways customized to different parts of the data analysis. For Table S1, Figure 1, and Figure S1, p-values were adjusted globally across pairs of comparisons, LB vs. WL and LRFR vs. WL in each genotype separately. For Table S5, Figure 3, and Figure S3, p-values were adjusted globally across three comparisons between different genotypes while keeping the light condition (WL) unchanged: *pif457* vs Col-0, *yuc2589* vs Col-0 and *smt2-1* vs Col-0. For Table S3, Table S4, Figure 2, Figure S2, Figure 3, and Figure S3, F-tests were used that yielded a single p-value for three comparisons, and post-hoc testing as implemented the R package limma was then applied to identify significantly regulated genes per comparison. For Figure S5, the same F-test is applied for two comparisons.

### Gene set enrichment analysis for Gene Ontology

Gene set enrichment analysis were conducted with ShinyGO v0.61:Gene Ontology Enrichment Analysis + more (http://bioinformatics.sdstate.edu/go/) (Ge et al., 2020) in *Arabidopsis thaliana* using a P-value cutoff (FDR) 0.05 and 500 most significant terms to show. The networks of enriched GO categories were visualized with R software (https://www.r-project.org/) using “visNetwork” and “igraph” libraries. Two terms (nodes) were connected if they share 20% or more genes. The size of the nodes indicates fold change for each term. We highlighted selected terms for each organ and light condition that we could easily relate to growth regulation (Fig 1D, S1C). The highlighted terms are not necessarily the most significant ones (full lists are available in Table S2 and as interactive versions).

### Statistical Motif Analysis in Promoter or Upstream Gene Sequences

Motif enrichment analyses were conducted with TAIR’s Motif Analysis tool (https://www.arabidopsis.org/tools/bulk/motiffinder/index.jsp) using 1kb upstream sequences.

### Other statistical analyses and data representation

For all the phenotypic analyses of hypocotyl elongation and the quantification of DII-signal, we performed two-way analysis of variance (ANOVA) (aov) and computed Tukey’s Honest Significance Differences (HSD) test (“agricolae” package) with default parameters using R software (https://www.r-project.org/). For phenotypic analysis of treatments to fenpropimorph and picloram, we performed two-way analysis of variance (ANOVA) (aov) and represented the significance as genotype*drug interaction in the given light conditions. For the comparisons including qPCR, ChIP-qPCR, sterol measurements, and lipidomics analysis, autophagic flux assays, we performed Student’s T-test. For lipidomics analysis, we further applied a BH correction for p values. We used binomial distribution for the PIF4 target enrichment, promoter motif, and to determine the significance of PIFs and/or YUCs dependence of enriched GO terms in given light conditions (Table S3, S4, S5).

In boxplots, the horizontal bar represents the median; boxes extend from the 25th to the 75th percentile, whiskers extend to show data range. In bar charts data are shown as the means ± standard deviation (SD). Autophagic flux assays are given as individual values from 4 biological replicates (average of 2 technical replicates), the horizontal lines indicate median, error bars extend to show data range. For the fenpropimorph and picloram treatments, the data show the means ± SD with a simple linear regression line connecting data points for the given light condition and the genotype. Asterisks (*) and different letters in the graphs indicate significant difference as defined by the statistical methods described above (P < 0.05).

## Notes

### Competing Interest Statement

The authors have declared no competing interest.

