## Supplemental Figures for "Reduced light access promotes hypocotyl growth via autophagy-mediated recycling"

**Figure S1.** LB and LRFR induce distinct transcriptional changes in cotyledons.
**Figure S2.** PIFs & YUCs are required for hormone-associated transcriptional changes in LRFR in cotyledons.
**Figure S3.** PIFs & YUCs are required for basal expression of many growth- and hormone-associated genes in WL in cotyledons.
**Figure S4.** SMT2 is locally required for LRFR-induced hypocotyl elongation.
**Figure S5.** Phenotyping evaluation of *cotyledon vasculature pattern* (*cvp*) mutants and *smt2-1* transcriptome summary.
**Figure S6.** Sterol composition of *B. rapa* hypocotyls do not change in LRFR
**Figure S7.** LB induces autophagy.

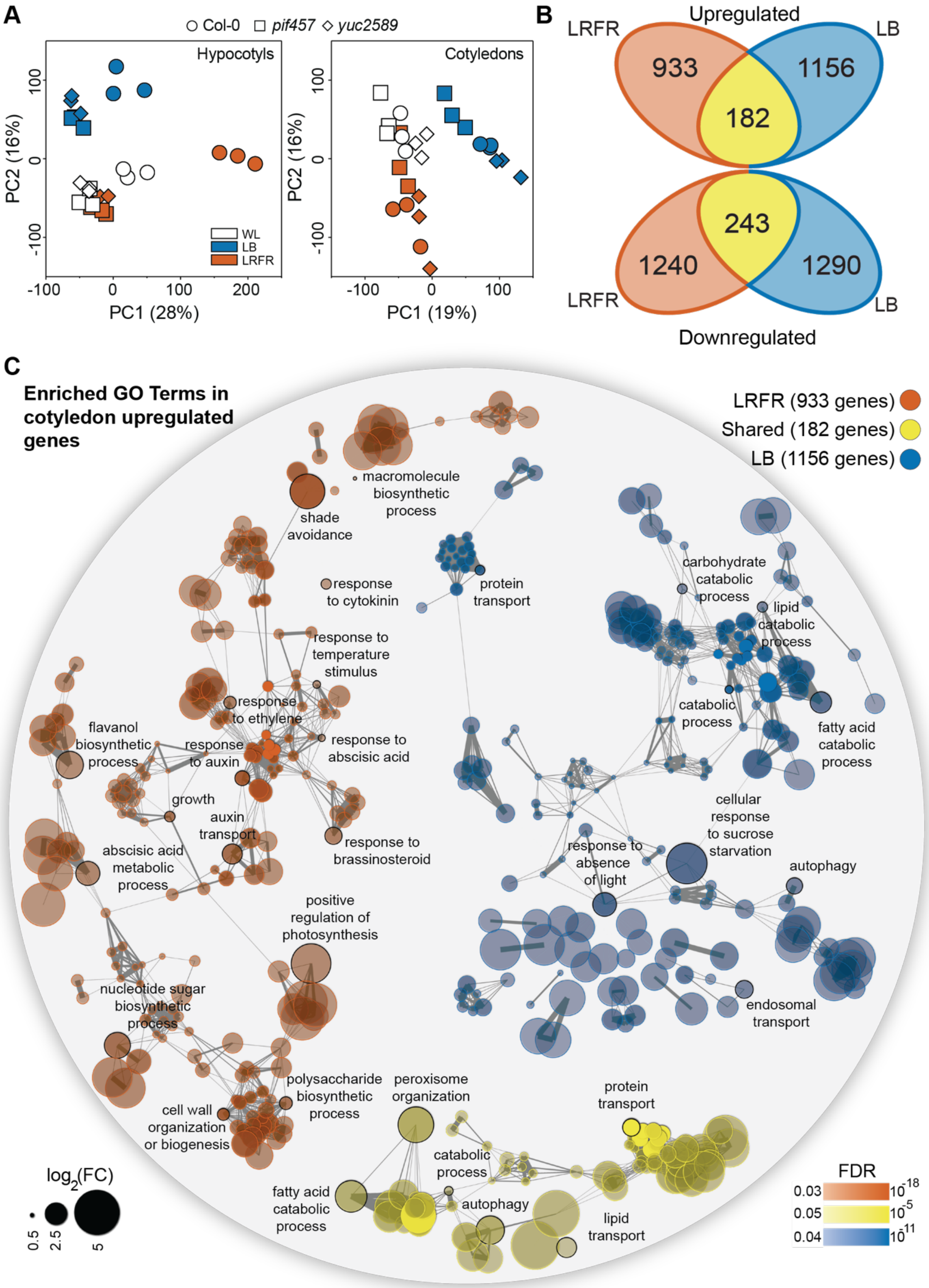


**Figure S1. LB and LRFR induce distinct transcriptional changes in cotyledons.**

(A) Principle component (PC) analysis of hypocotyl and cotyledon transcriptomes. PC1 and PC2 of each biological replicate (n = 3) are graphically visualized. (B) Number of up- and downregulated genes in Col-0 cotyledons in the indicated light conditions (FDR<0.05, T-test with BH correction). (C) GO term enrichment analysis in the cotyledon upregulated gene lists. Each node indicates a significantly enriched GO term

(FDR<0.05). Two terms (nodes) are connected if they share 20% or more genes. The line thickness increases with the increased number of shared gene sets between two terms. Only selected GO terms (black circles) are annotated. The full list of enriched GO terms is in Table S2, the interactive version of (C) is available here. See also Figure 1.

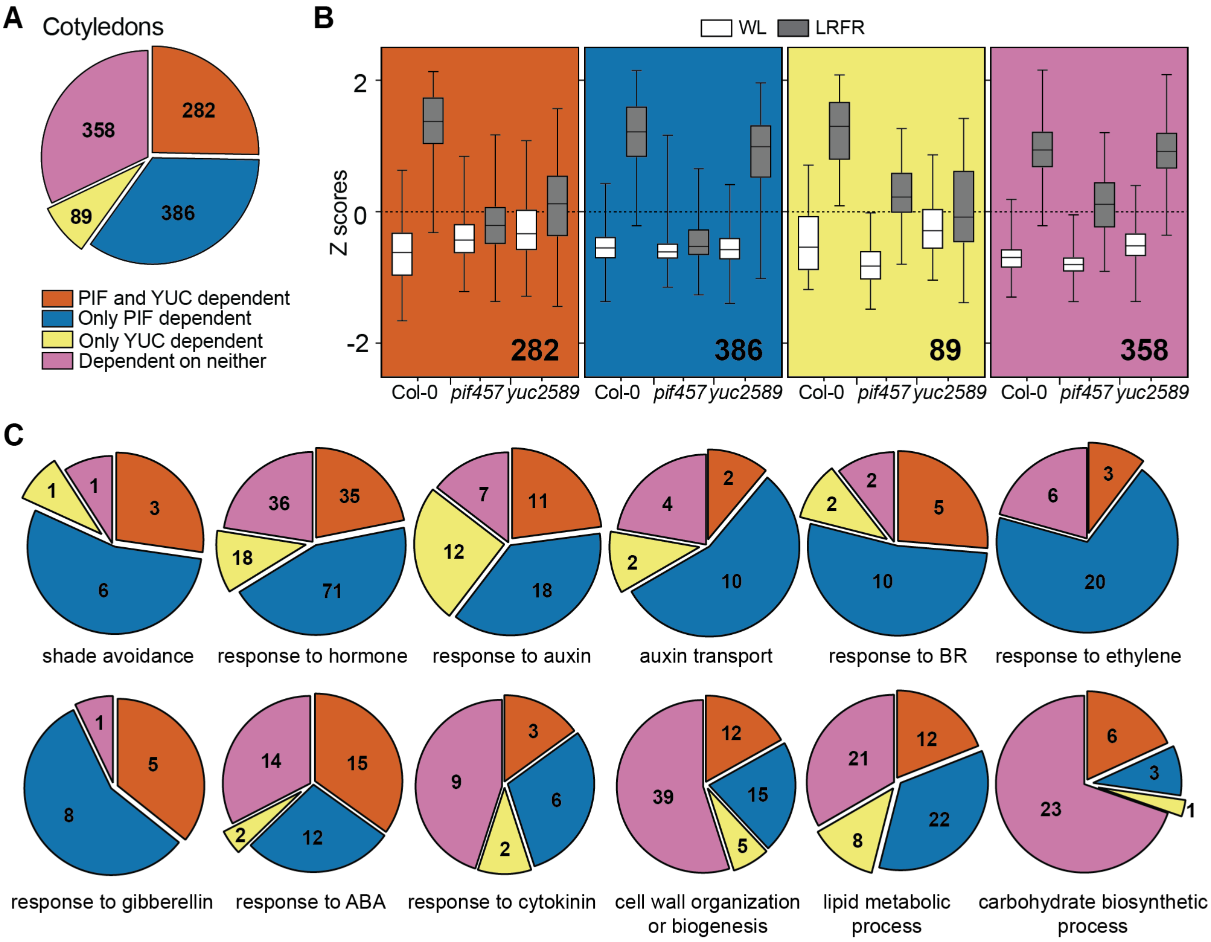


**Figure S2. PIFs & YUCs are required for hormone-associated transcriptional changes in LRFR in cotyledons.**

(A) The distribution of cotyledon-induced genes in LRFR according to the dependence on PIFs and YUCs using the comparison of Col-0, *pif457* and *yuc2589* transcriptomes (FDR<0.05, F-test with post-hoc test). (B) Distributions of Z- scores computed from replicates averages for categories shown in (A). The horizontal bar represents the median; boxes extend from the 25th to the 75th percentile, whiskers extend to show the data range. (C) The distribution of cotyledon-induced genes according to the dependence on PIFs and YUCs in each of the selected significantly enriched GO terms. Numbers indicate significantly regulated genes in the given categories and/or GO terms. The full list of misregulated genes and enriched GO terms are given in Table S3. See also Figure S2.

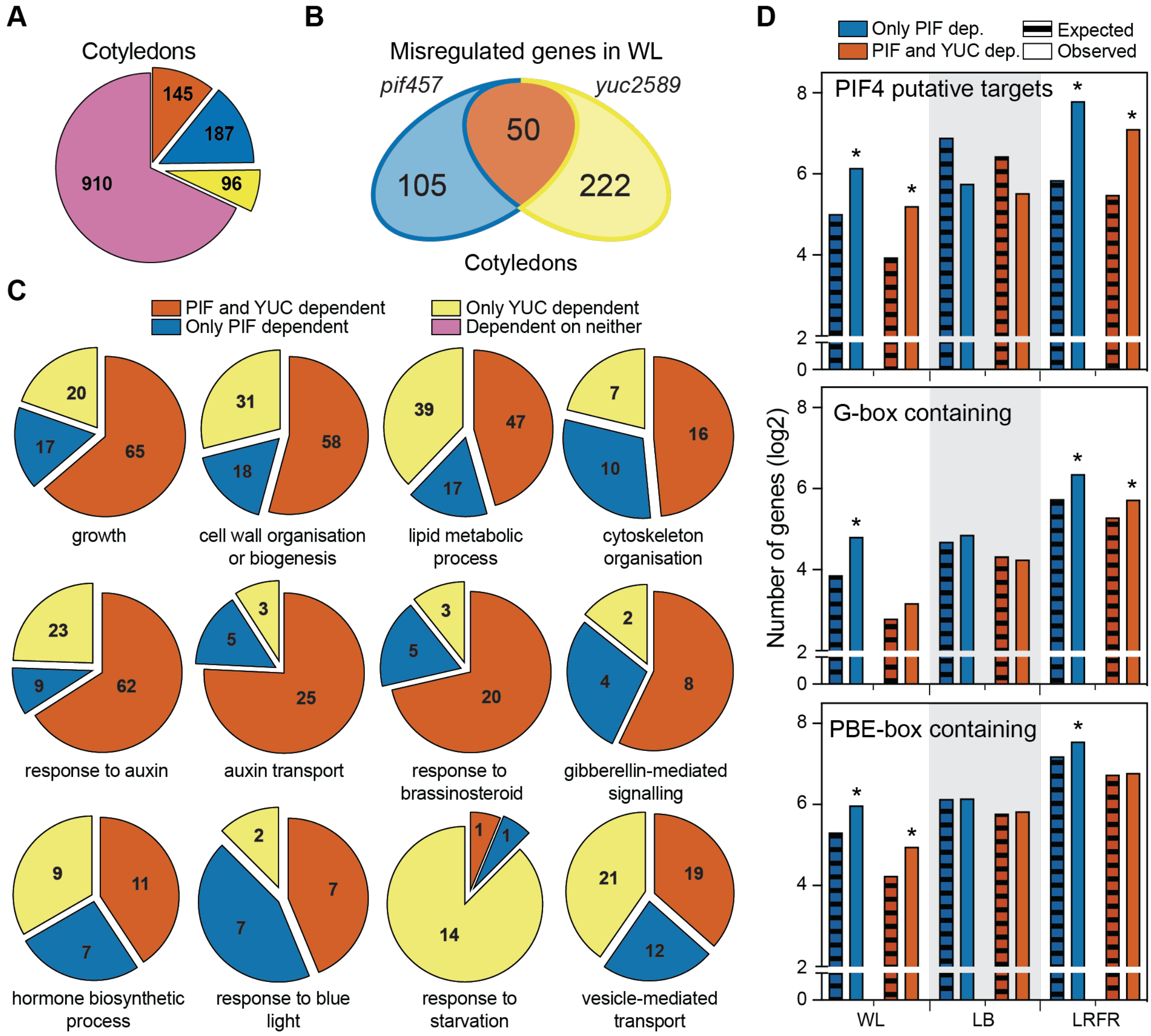


**Figure S3. PIFs & YUCs are required for basal expression of many growth- and hormone- associated genes in WL in cotyledons.**

(A) The distribution of cotyledon-induced genes in LB according to the dependence on PIFs and YUCs using the comparison of Col-0, *pif457* and *yuc2589* transcriptomes (FDR<0.05, F-test with post-hoc test). (B) Number of misregulated genes *pif457* and *yuc2589* hypocotyls compared to Col-0 in WL (FDR<0.05, T-test with BH correction). (C) The distribution of misregulated genes in *pif457* and *yuc2589* hypocotyls (as grouped in Fig. 3C) in each of the selected significantly enriched GO terms. The full list of enriched GO terms is given in Table S5. (D) Comparison of PIF dependent genes in cotyledons with PIF4 putative targets (as listed in (Pedmale et al., 2016)), promoters (1 kb upstream) containing G-box (CACGTG) or PBE-box (CATGTG). Asterisks (*) indicate the statistically significant overrepresentation compared to expected ($P < 0.05$, Binomial distribution). Numbers indicate significantly regulated genes in the given categories and/or GO terms. The full list of misregulated genes, enriched GO terms, and enriched motifs are given in Table S4, S5, and S6. See also Figure 3.

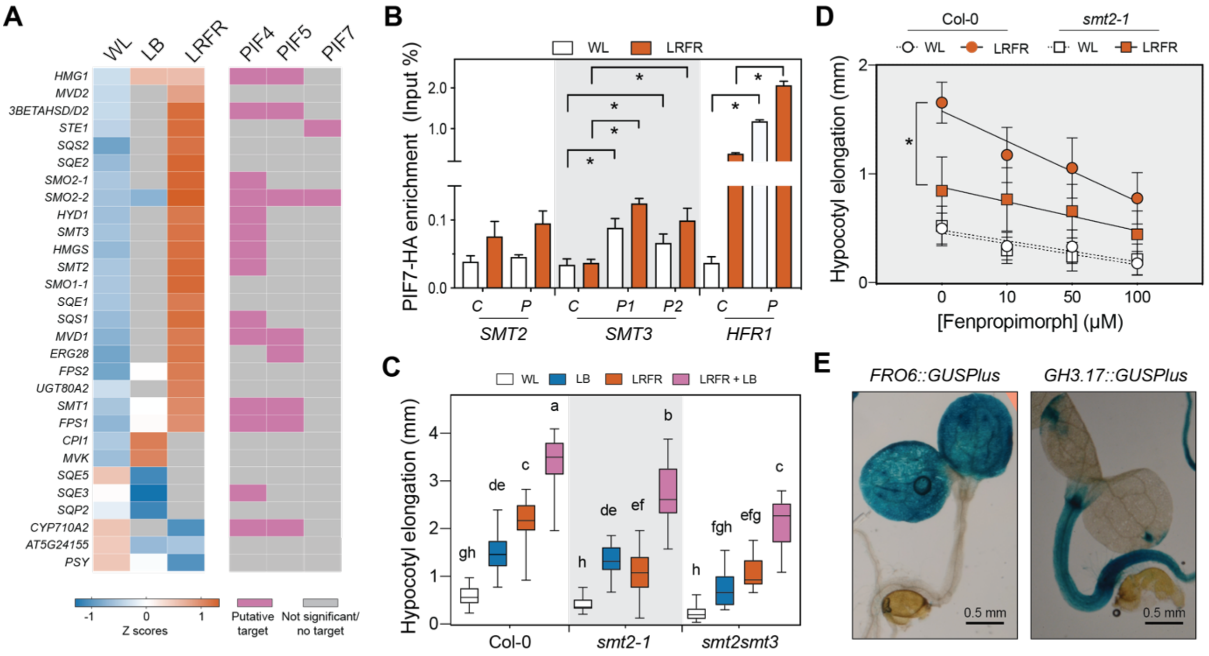


**Figure S4. SMT2 is locally required for LRFR-induced hypocotyl elongation.**

(A) The heatmap showing average Z-scores of genes annotated in sterol biosynthesis (GO: 0016126) in RNA-seq and putative binding of PIF4, PIF5 and PIF7 (ChIP-seq data from (Chung et al., 2020, Pedmale et al., 2016)). A gene is shown if it is significantly regulated at least in one treatment (LB or LRFR). (B) PIF7-HA binding to the promoter of the indicated genes. PIF7-HA enrichment is quantified by qPCR and presented as IP/Input (n=3, technical). Control – C, Peak – P. (C, D) Hypocotyl elongation of indicated genotypes in the indicated conditions (n>12). (C) The horizontal bar represents the median; boxes extend from the 25th to the 75th percentile, whiskers extend to show the data range. (B, D) Data are means ± SD. Different letters (C, two-way ANOVA with Tukey's HSD test) and asterisks (*) (B, T-test) (D, two-way ANOVA) indicate significant difference ($P < 0.05$) compared to control (B), and between genotypes in given light condition (D). (E) GUS staining of the indicated *promoter::GUSPlus* lines. See also Figure 4.

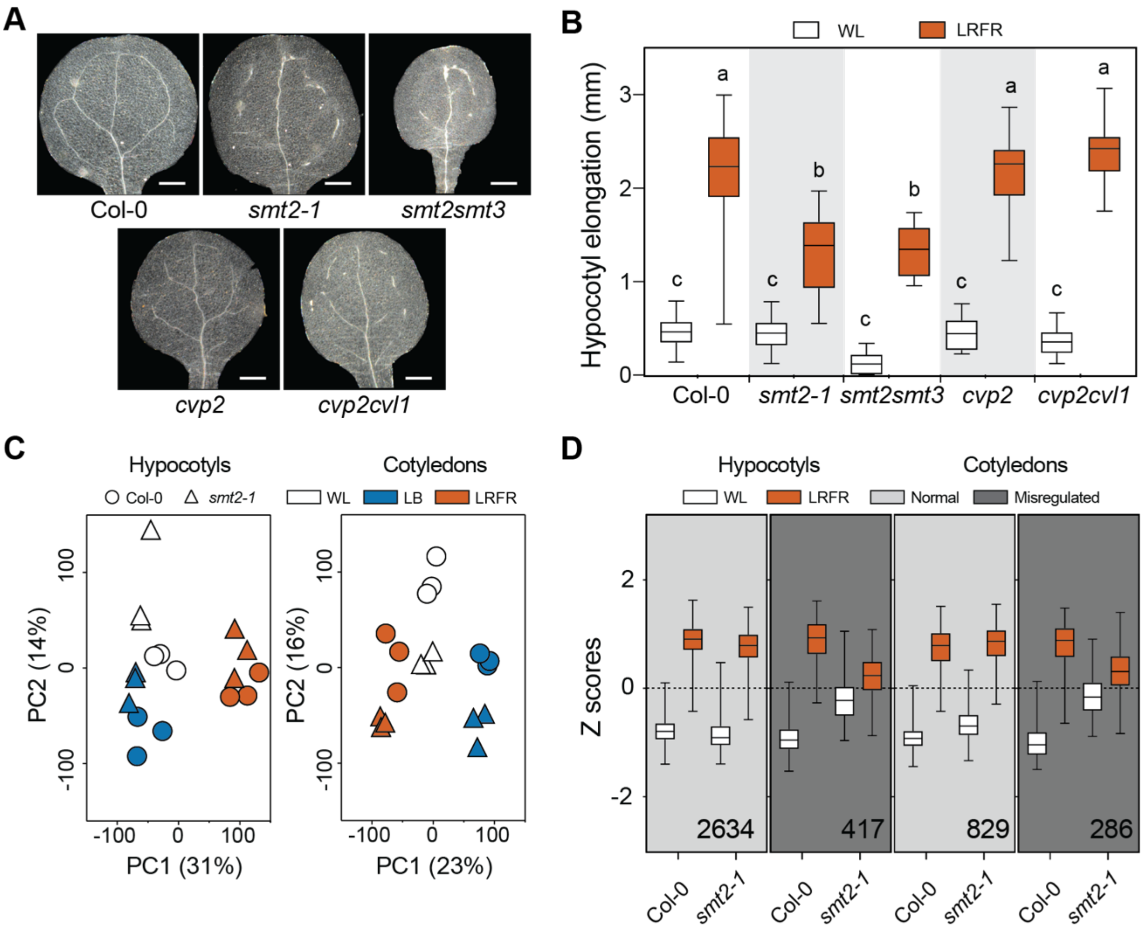


**Figure S5. Phenotyping evaluation of *cotyledon vasculature pattern* (*cvp*) mutants and *smt2-1* transcriptome summary.**

(A) Representative images of cotyledon vasculature phenotype of indicated genotypes. White bars equal to 200 µm. (B) Hypocotyl elongation of indicated genotypes. Different letters indicate significant difference ($P < 0.05$, $n > 12$, two-way ANOVA with Tukey's HSD test). (C) PCA of Col-0 and *smt2-1* transcriptomes. PC1 and PC2 of each biological replicate ($n = 3$) are graphically visualized. (D) Distributions of Z- scores computed from replicates averages for LRFR-induced genes that are grouped comparing Col-0 and *smt2-1* transcriptomes (FDR<0.05). 'Normal' and 'Misregulated' refer to expression in *smt2-1*. The numbers indicate significantly regulated genes in each group (FDR<0.05, F-test with post-hoc test). (B, D) The horizontal bar represents the median; boxes extend from the 25th to the 75th percentile, whiskers extend to show the data range. The full list of misregulated genes is given in Table S7. See also Figure 5.

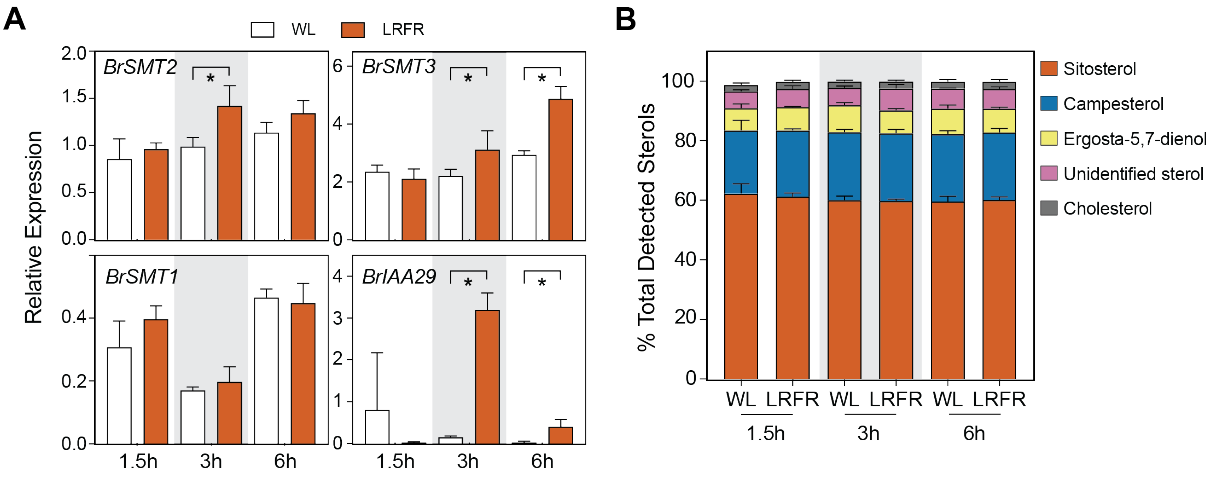


**Figure S6. Sterol composition of *B. rapa* hypocotyls do not change in LRFR**

(A) Relative expression of the indicated genes in 5-d-old *B. rapa* hypocotyls. Gene expression values were calculated as fold induction relative to *BrPP2A*. n = 4 (biological) with three technical replicas for each RNA sample. Asterisks (*) indicate $P < 0.05$ (T-test). (B) Sterol composition of 5-d-old *B. rapa* hypocotyls (n=5, biological). (A, B) Data are means ± SD. See also Figure 6.

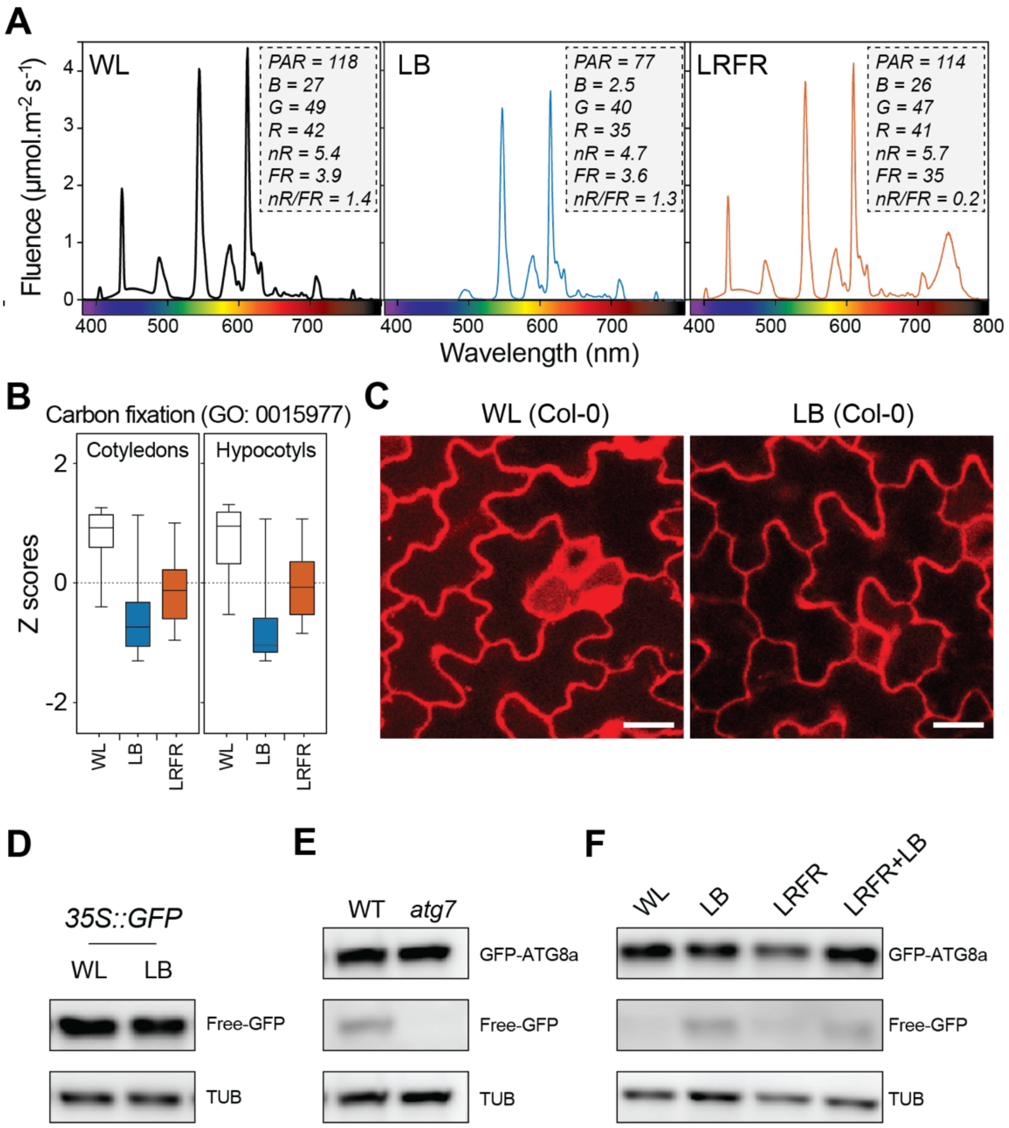


**Figure S7. LB induces autophagy.**

(A) Light conditions used in this study. Photosynthetically active radiation (PAR, 400-700 nm), Blue (B, 400-500 nm), Green (G, 500-600 nm), Red (600-700 nm), Narrow R (nR, 640-700) and far-red (FR, 700-760 nm). (B) Distributions of Z- scores computed from replicates averages for genes listed in carbon fixation GO term in Col-0 seedlings. The horizontal bar represents the median; boxes extend from the 25th to the 75th percentile, whiskers extend to show the data range. (C) Cotyledon pavement cells of *UBQ10::mCherry-ATG8e* in Col-0 kept in WL or treated with LB in the absence of ConA for 8h. White bars equal to 20 µm. (D) GFP levels are detected in *35S::GFP* line in 8h of WL and LB without CA. GFP-ATG8a and Free-GFP levels are detected in (E) WT and *atg7-2* seedlings treated with 8h of LB in the presence of 0.5 µM ConA and (F) in WT seedlings treated as in (Fig. 7F) without ConA. TUB was used as a loading control (D, E, F). The full list of genes in Carbon fixation GO term is given in Table S10. See also Figure 7.
